# Dynamic Bacterial Growth Modulation in Structurally Distinct and Functionally Tuneable Agarose Hydrogels

**DOI:** 10.1101/2025.06.28.662093

**Authors:** Andrea Dsouza, Dylan Taylor, Christopher Parmenter, Rachel A. Hand, Julia Brettschneider, Meera Unnikrishnan, Chrystala Constantinidou, Jérôme Charmet

**Affiliations:** Division of Biomedical Sciences, Warwick Medical School, The University of Warwick, Coventry, CV4 7AL, United Kingdom; Big Data Institute, Li Ka Shing Centre for Health Information and Discovery, Nuffield Department of Population Health, University of Oxford, UK; Nanoscale and Microscale Research Centre, The University of Nottingham, Nottingham, NG7 2RD, United Kingdom; Department of Chemistry, The University of Warwick, CV4 7AL, United Kingdom; Department of Statistics, The University of Warwick, Coventry, CV4 7AL, United Kingdom; Bioinformatics Research Technology Platform, The University of Warwick, CV4 7AL, United Kingdom; School of Engineering - HE-Arc Ingénierie, HES-SO University of Applied Sciences Western Switzerland, 2000 Neuchâtel, Switzerland; School of Precision and Biomedical Engineering, University of Bern, Güterstrasse 24/26, 3008 Bern, Switzerland

**Keywords:** Hydroxyethylated agarose hydrogels, hydrogel stiffness, hydrogel water loss, cytochrome C, electrostatic interactions, bacterial growth, bacterial inhibition

## Abstract

Bacterial adaptability to diverse environments drives infection, persistence, and antibiotic resistance. Although hydrogels are increasingly used to model such conditions, the factors governing hydrogel-dependent bacterial growth is complex. Here, we focus on agarose hydrogels and investigate how their material properties influence bacterial proliferation. Using two agarose types – hydroxyethyl substituted and unsubstituted – at varying concentrations, we tested four bacterial species (*E. coli, P. fluorescens, S. aureus, B. subtilis*) across five nutrient media yielding 120 conditions. Growth consistently decreased with increasing hydrogel stiffness and water loss in unsubstituted and substituted agarose hydrogels, regardless of species. Media effects were largely due to their impact on hydrogel properties rather than nutrient content. Furthermore, electrostatic repulsion between Gram positive bacteria and anionic unsubstituted agarose suppressed growth in high concentration gels. These findings demonstrate that bacterial growth in agarose systems is primarily shaped by gel mechanics and surface interactions, informing the design of infection models and antibacterial materials.

## Introduction

Bacteria have acquired an extraordinary ability to adapt and thrive in various microenvironments due to survival mechanisms including formation of complex biofilms^1,2^, exchange of genetic material by horizontal gene transfer^3–6^, and antimicrobial resistance^7,8^. Consequently, the incidence of bacterial infections, particularly antibiotic-resistant infections, is increasing faster than ever and has claimed approximately 4.71 million deaths globally in 2021 alone^9^. Life-threatening infections are also a major driver of significant healthcare and economic costs, and results in socio-economic devastation and heightened vulnerability in low-to-middle income countries^10–13^. The World Health Organization has declared antimicrobial resistance as one of the top global public health and development threats^14,15^, highlighting the urgent need for new diagnostics and better treatments.

Due to their unique characteristics including porosity, elasticity, biocompatibility, and ease of customization, hydrogels have been extensively designed for exploring bacterial interactions, antibacterial formulations, wound dressings, and smart biosensors for bacterial detection^16–21^. Hydrogels are considered one of the most promising *in vitro* models for studying bacterial growth and anti-bacterial activities^16^. In particular, agarose hydrogels are valued in microbiology for their chemical inertness, structural simplicity, and widespread use in culturing and visualizing bacterial cells.

Despite growing interest in using hydrogels to understand and control bacterial behaviour, the fundamental properties that govern bacterial growth within these matrices remain poorly understood. Properties such as stiffness, porosity, hydrophilicity, and nutrient content have been implicated^22-25^, but findings, particularly concerning stiffness have been inconsistent. For instance, some studies report increased bacterial growth with increased stiffness, while others observe the opposite. In agarose-based systems, Guegan et al. found that Gram-negative *Pseudoalteromonas sp*. adhered more effectively to stiffer gels, whereas *Bacillus subtilis* showed no preference^26^. Similarly, increased mechanical stress in stiff agarose has been shown to enhance *Pseudomonas aeruginosa* accumulation^29^. However, contradictory results in other materials, such as reduced *Staphylococcus aureus* adhesion on stiffer polyacrylamide substrates, suggest that hydrogel chemistry, not stiffness alone, governs bacterial response^31^. Furthermore, bacterial envelope characteristics, particularly surface charge and production of extracellular polymeric substances are often underrepresented in these analyses, despite their key role in hydrogel-bacteria interactions^16,32,33^.

Given these complexities, our study focuses specifically on agarose hydrogels, to decouple and systematically investigate the physicochemical parameters that regulate bacterial growth. We used two structurally distinct agarose derivatives: (i) unsubstituted agarose (US), containing only hydroxyl (-OH) groups, and (ii) substituted agarose (S), containing hydroxyethyl (-C_2_H_5_OH) groups in addition to hydroxyls. Though derived from the same polysaccharide backbone of D-galactose and 3,6-anhydro-L-galactopyranose, these modifications confer differing mechanical and hydration properties to the gels.

To evaluate how these structural differences influence bacterial behaviour, we examined the growth of four representative bacterial species: *Escherichia coli* and *Pseudomonas fluorescens* (Gram-negative), and *Staphylococcus aureus* and *Bacillus subtilis* (Gram-positive) across both hydrogel types. We further varied hydrogel concentration (0.2%, 0.5%, and 1%) and encapsulated each with five commonly used bacterial nutrient media including Nutrient Broth (NB), Lysogeny Broth (LB), Tryptic Soy Broth (TSB), and two formulations of Mueller Hinton Broth (M1 and M2), resulting in 120 experimental conditions.

Our results reveal that bacterial growth in agarose hydrogels is predominantly governed by stiffness and water loss, with growth consistently increasing in softer, more hydrated hydrogels across all species. Substituted hydrogels, which exhibit lower stiffness and reduced water loss, supported greater bacterial growth than unsubstituted counterparts. Interestingly, the effect of nutrient media was mediated more through its impact on hydrogel physical properties than direct nutrient content. Additionally, we observed that electrostatic repulsion between negatively charged bacteria and the anionic unsubstituted hydrogels inhibited growth in certain conditions, particularly at higher gel concentrations. Collectively, these findings highlight the critical role of hydrogel physicochemical properties and bacterial envelope characteristics in modulating bacterial growth behaviour within agarose-based systems. While our study is limited to agarose hydrogels, the observed species-and material-dependent variability suggests that generalizations based solely on bulk mechanical properties may be insufficient. These results underscore the need for a material-specific framework to guide the rational design of hydrogels as antibacterial biomaterials, infection models, and platforms for studying microbe–material interactions.

## Results

### Low hydrogel concentrations promote increased bacterial growth

To investigate the effect of hydrogel concentration, we first studied bacterial growth separately in US and S agarose hydrogels. These hydrogels were prepared at concentrations of 0.2%, 0.5%, and 1%, each encapsulating a defined nutrient medium (NB, LB, TSB, M1, and M2). Bacterial cultures were inoculated at the centre of the gels and incubated at 37°C for 18 hours. Growth areas (Figure S2-S9) were quantified using Fiji software (Figure S1).

Our results demonstrate that the growth of uropathogenic *E. coli, P. fluorescens, S. aureus, and B. subtilis* increases progressively as hydrogel concentration decreases (Figure 2). Notably, reductions from 1% to 0.2% and from 0.5% to 0.2% led to 82.5% and 55% increases in bacterial growth, respectively, across both US and S hydrogels. Even a modest concentration decrease from 1% to 0.5% results in a 27.5% growth increase, indicating that bacterial growth is highly sensitive to small changes in hydrogel concentration. These findings demonstrate a strong positive effect of lower hydrogel concentrations on bacterial growth.

**Figure 1.**
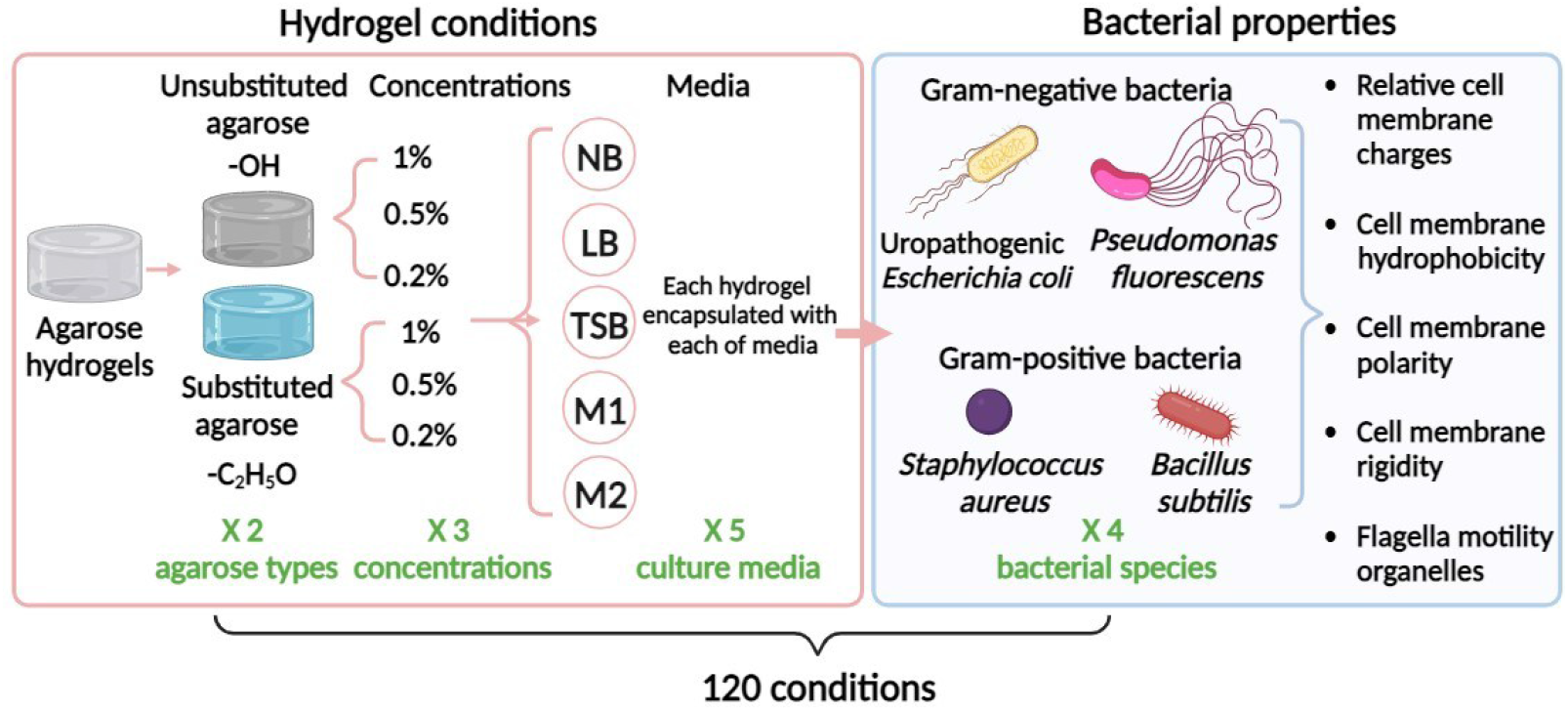
Schematic illustration of the multiparametric study involving hydrogel conditions and bacterial properties. 2 types of agarose hydrogels namely, unsubstituted and substituted hydrogels, each at 3 different concentrations of 1%, 0.5%, and 0.2% encapsulated with 5 different nutrient media including NB, LB, TSB, M1, and M2 were included for assessment of bacterial growth. The growth of 4 different bacterial species including *E. coli* and *P. fluorescens* (Gram-negative), and *S. aureus* and *B. subtilis* (Gram-positive) bacterial species were studied. In total the study included 120 conditions.

**Figure 2.**
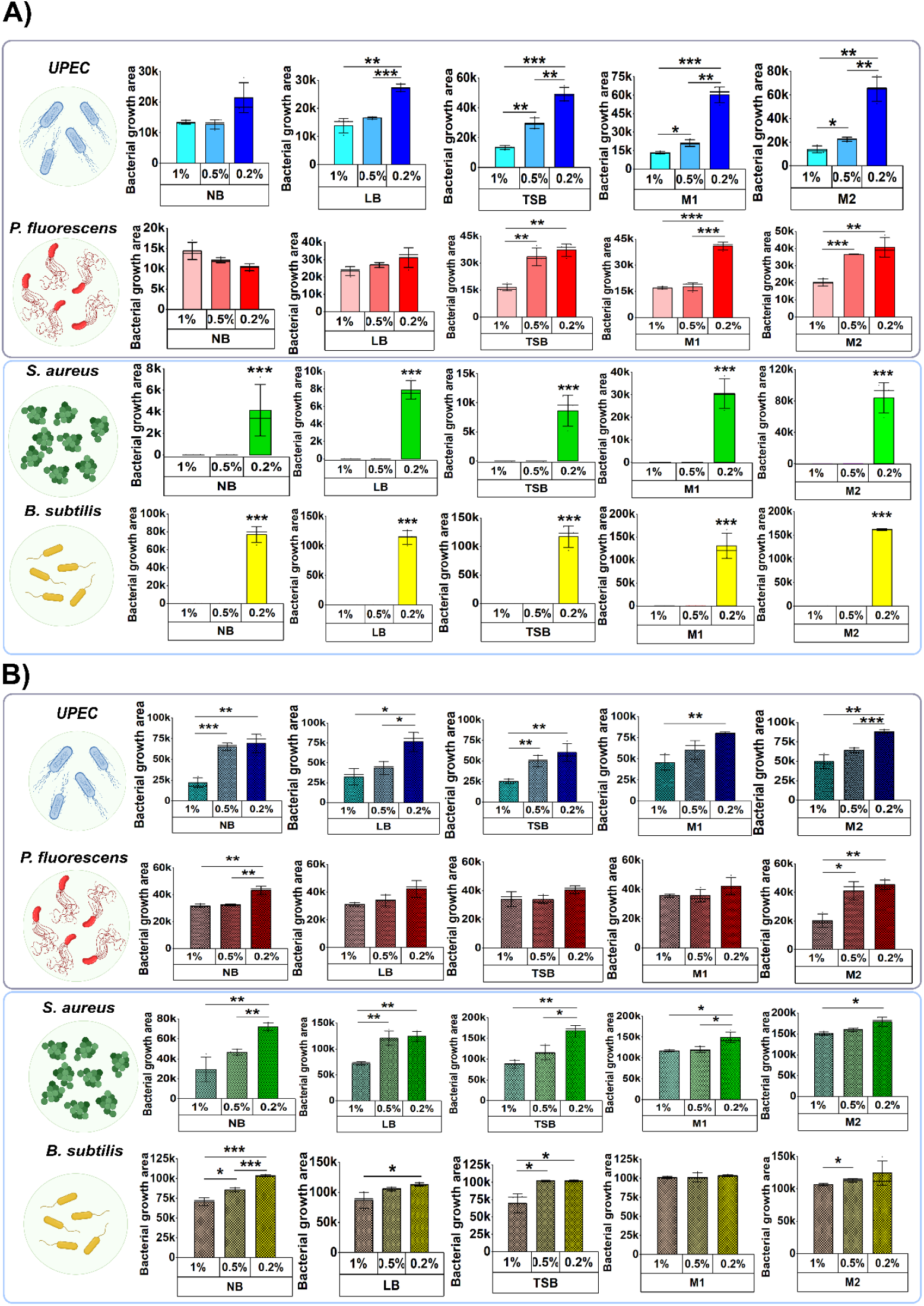
Quantitative growth of (i) uropathogenic *E. coli*, (ii) *P. fluorescens*, (iii) *S. aureus*, and (iv) *B. subtilis* in NB, LB, TSB, M1, and M2 – encapsulated unsubstituted **(A)** and substituted **(B)** agarose hydrogels, each at 1%, 0.5%, and 0.2% hydrogel concentrations. Data are presented as mean ± standard deviation (SD) from n=3 independent replicates. Statistical significance was assessed using ANOVA. \**P* < 0.05, \*\**P* < 0.01, and \*\*\**P* < 0.001.

To explore structural differences underlying the observed growth patterns, we performed cryo-SEM imaging of US and S hydrogels at varying concentrations (Figure S10). In US hydrogels, higher agarose concentrations (1%) showed smaller, denser pores, while lower concentrations (0.5%) exhibited visibly larger pores. S hydrogels displayed a distinct thread-like network, with 1% gels forming more compact structures than 0.5%. These morphological differences suggest that lower hydrogel concentrations support greater porosity, potentially enhancing nutrient diffusion and facilitating bacterial proliferation.

### Hydrogel stiffness scales with polymer concentration

To investigate the physical basis for the observed concentration-dependent increase in bacterial growth, we assessed how polymer concentration influences the mechanical properties of the hydrogels. Quantitative rheological measurements showed a marked decrease in storage modulus (G′) with decreasing agarose concentration, consistent across both US (Figure 3B) and S (Figure 3C) hydrogels. Specifically, stiffness decreased from 3825 ± 630 Pa (US) and 1479 ± 270 Pa (S) at 1%, to 1130 ± 300 Pa (US) and 245 ± 22 Pa (S) at 0.5%, and further to 240 ± 24 Pa (US) and 156 ± 46 Pa (S) at 0.2%. The amplitude and frequency sweep curves of hydrogels are provided in Figures S3-S12. The reduction in stiffness aligns with the increased bacterial growth observed at lower concentrations, suggesting that softer hydrogel networks provide a more permissive environment for proliferation. Amplitude and frequency sweeps are provided in Figure S11-S20.

**Figure 3.**
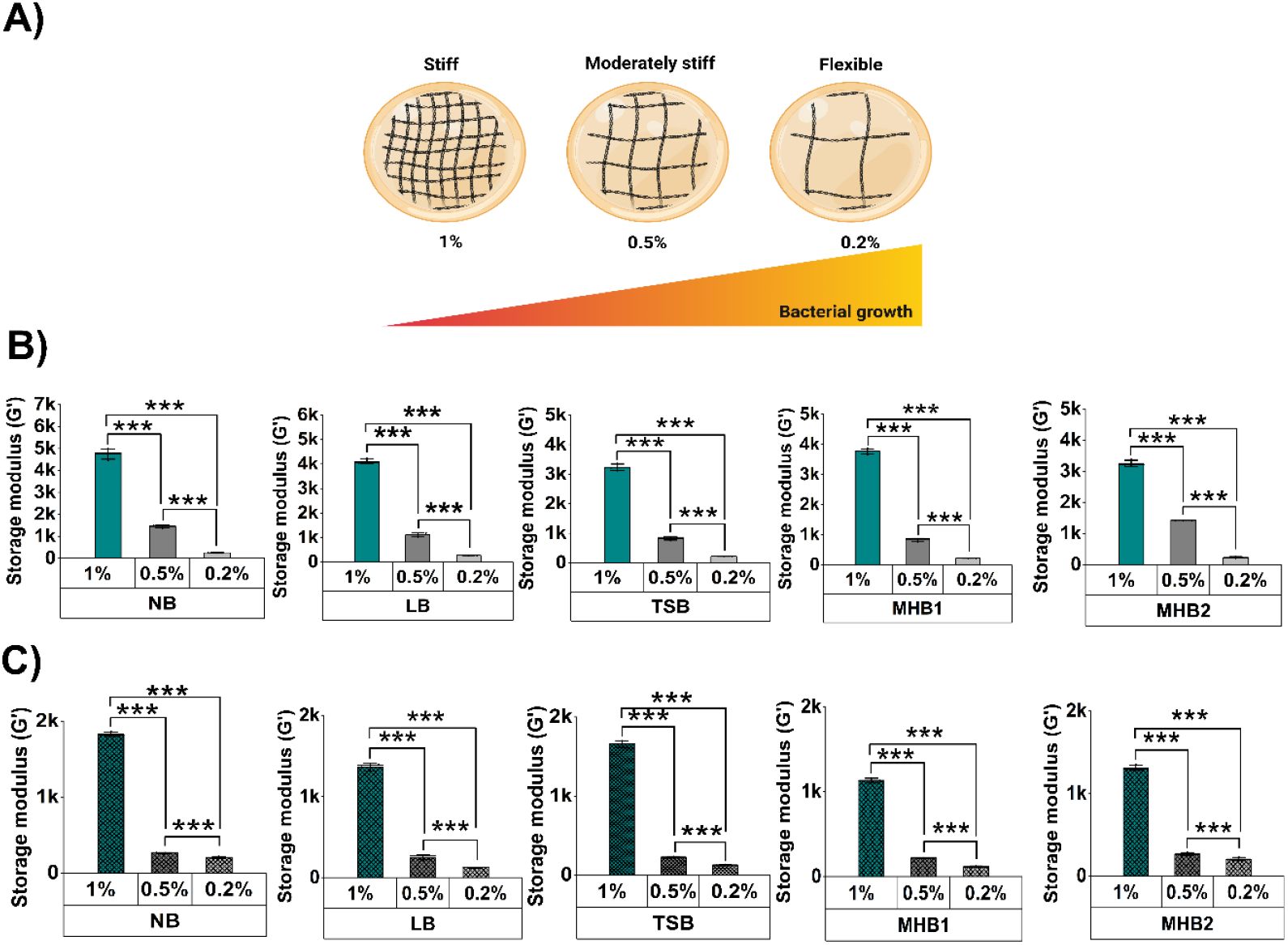
**A)** Schematic showing decreasing agarose concentrations (1% → 0.5% → 0.2%) with varying crosslinking and stiffness. Storage modulus (G’) of **(B)** US and **(C)** S hydrogels with NB, LB, TSB, M1, and M2 at varying concentrations. Data are presented as mean ± standard deviation (SD) from n=3 independent replicates. Statistical significance was assessed using ANOVA. *P* value: \**P* < 0.05, \*\**P* < 0.01, \*\*\**P* < 0.001.

### Lower polymer content preserves hydrogel water retention

Alongside stiffness evaluation, we also determined the water retention capacity of our hydrogels. Studies have suggested that the water content of hydrogels may influence bacterial growth behaviour^29,41,42^; however, this aspect remains understudied. Indeed, the hydrophilicity of the hydrogel play a key role in regulating water retention, nutrient availability, and diffusion within the hydrogel network, thereby influencing bacterial colonization. Determination of water retention is even more important in cases where hydrogels are incubated at 37°C for 18 hours, as water evaporates quickly during this time, interfering with bacterial growth. To assess this effect, we measured the weights of hydrogels before and after incubation. Interestingly, we identified the highest and statistically significant water loss in the stiffest 1% hydrogels compared to intermediately stiff 0.5% and flexible 0.2% hydrogels, in both US (Figure 4B) and S (Figure 4C) hydrogels across all nutrient media encapsulated formulations.

**Figure 4.**
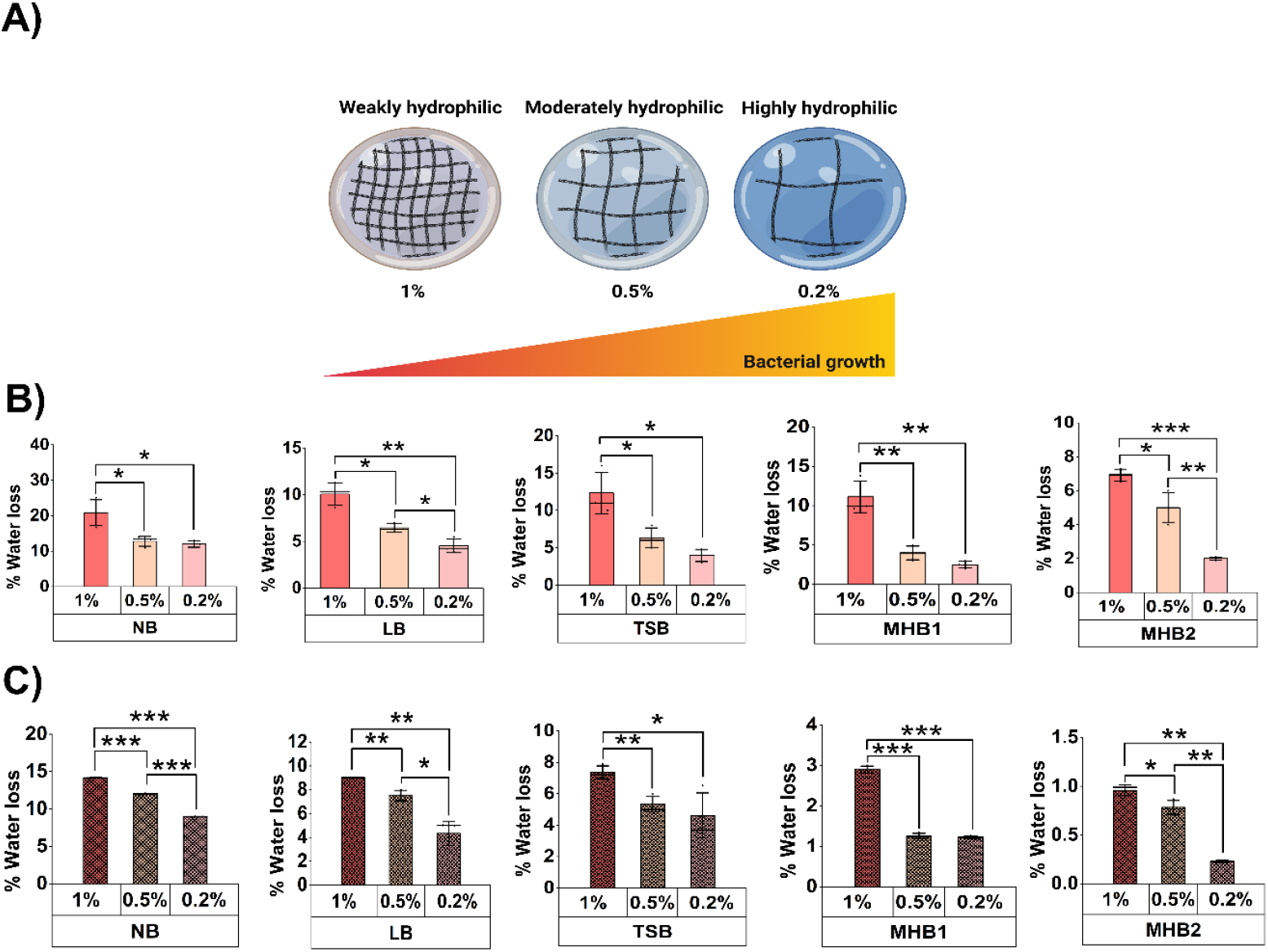
**A)** Schematic showing decreasing agarose concentrations (1% → 0.5% → 0.2%) with varying crosslinking and hydrophilicity. % Water loss of **(B)** US and **(C)** S hydrogels with NB, LB, TSB, M1, and M2 at varying concentrations. Data are presented as mean ± standard deviation (SD) from n=3 independent replicates. Statistical significance was assessed using ANOVA. *P* value: \**P* < 0.05, \*\**P* < 0.01, \*\*\**P* < 0.001.

### Combined effects of stiffness and water loss on bacterial growth

To provide a broader overview of the influence of hydrogel stiffness and water loss on bacterial growth, we analysed and combined the data into heatmaps (Figure S22). When considering US and S hydrogels separately, one observes increasing bacterial growth from 1% to 0.2% for both stiffness and water loss. When this data is combined into a single heatmap (with US on the left and S on the right), a non-linear pattern, as shown for *E*.*coli* in Figure 5A, becomes evident, with all values seeming to increase from left to right, with the exception of the third column (US 0.2%) that consistently exhibits higher values (for stiffness, water loss and growth) than the fourth column (S 1%). This clear repeating pattern hints at a strong correlation between bacterial growth, stiffness and water loss, independent from the hydrogel type. This observation is confirmed in Figure 5B that shows a consistent, and progressive growth pattern obtained after rearranging the heatmaps in order of decreasing stiffness and water loss.

**Figure 5.**
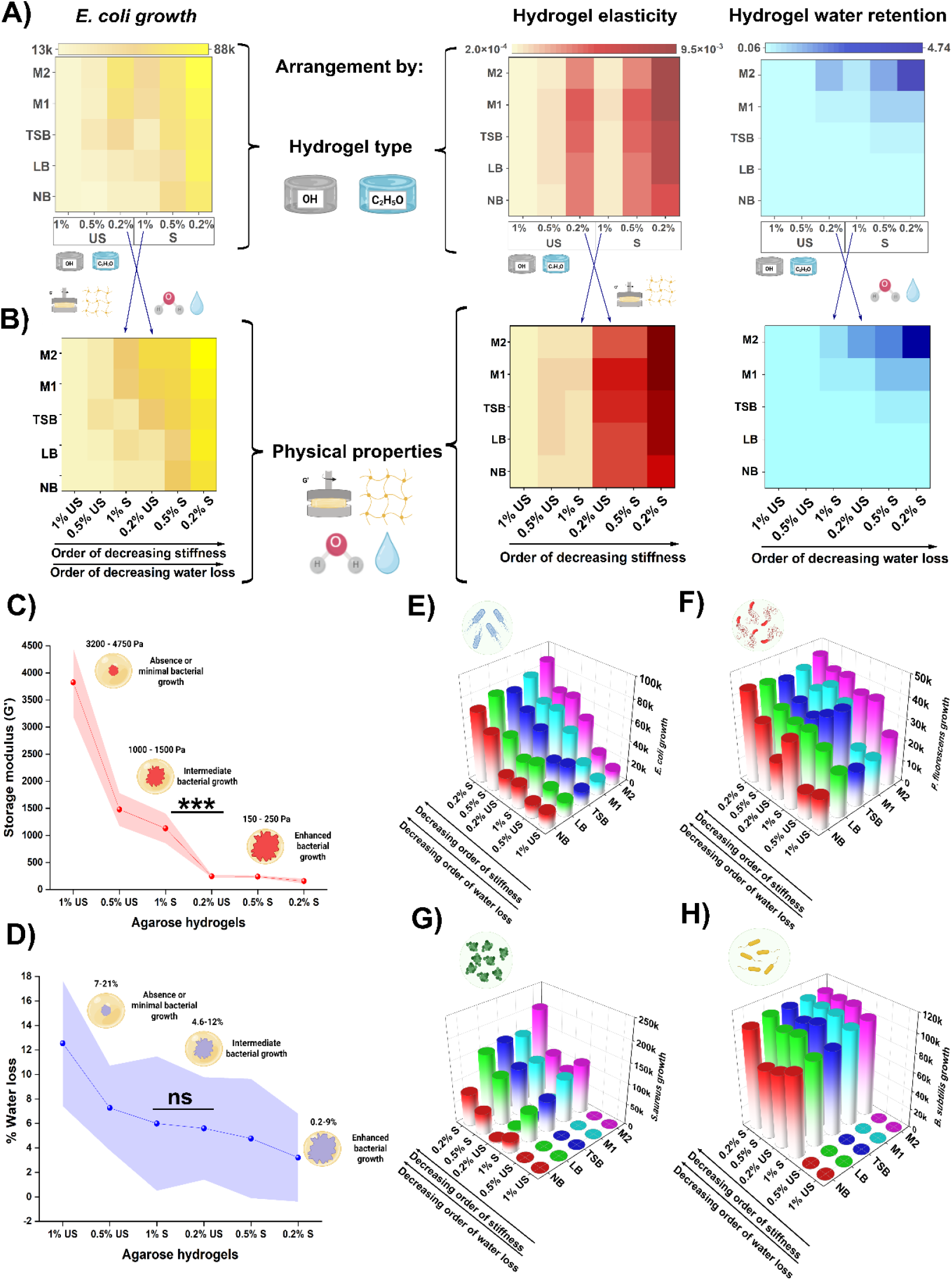
Heatmaps indicating the growth of *E. coli* according to **(A)** hydrogel type, hydrogel elasticity and water retention and **(B)** in order of decreasing hydrogel stiffness and water loss. **(C)** Storage modulus (G’) (***= *P* < 0.001) indicating increasing stiffness with US and S hydrogels combined: 0.2% S < 0.5% S < 0.2% US < 1% S < 0.5% US < 1% US. **(D)** % water in US and S hydrogels combined: 0.2% S < 0.5% S < (0.2% US = 1% S) < 0.5% US < 1% US. Maximum bacterial growth occurs in hydrogels with 150-250 Pa stiffness and 0.2-9% water loss. Growth of **(E)** *E. coli*, **(F)** *P. fluorescens*, **(G)** *S. aureus*, and **(H)** *B. subtilis* increases with decreasing hydrogel stiffness and water loss.

The above observation is not only limited to *E. coli* but is also observed in *P. fluorescens* growth (Figure S21). In contrast, *S. aureus* and *B. subtilis* showed more complex trends, involving electrostatic interactions (described below). Indeed, 3D heatmaps in Figure 5E-5H show the growth of all bacterial species arranged as function of stiffness and water loss irrespective of the hydrogel type. Similar trends are observed for all bacterial species; a negative association between both stiffness and water loss was seen with bacterial growth, as confirmed using Spearman’s correlation (Tables S23 and S24). We also note species-specific responses. While *E. coli* followed the trend, *P. fluorescens* showed slightly higher growth in 1% S than 0.2% US in some media, though differences were statistically insignificant (*P* > 0.05, Figure 5E and F). For *S. aureus* and *B. subtilis*, growth was inhibited in 1% and 0.5% US, but minimal growth was observed in 0.2% US (Figure 5G and H). Overall, these results demonstrate that decreased hydrogel stiffness and water loss enhance bacterial growth, with distinct growth patterns observed across different species.

### Encapsulated nutrient media impacts bacterial growth by altering hydrogel stiffness and water loss

Further supporting our findings that bacterial growth correlates with decreasing hydrogel stiffness and water loss, we observed a surprising effect of encapsulated nutrient media on bacterial growth. Growth of *E. coli, P. fluorescens, S. aureus*, and *B. subtilis* was tested in NB, LB, TSB, M1, and M2, both in-solution and in hydrogels. In-solution growth varied across media and species without a clear pattern (Figure 6A). When hydrogels (1%, 0.5%, and 0.2% US and S) were encapsulated with these media, 90% of the hydrogels showed significantly greater bacterial growth in M2 compared to NB. Heatmap analysis (Figure 6B) revealed that M2 encapsulation led to increased bacterial growth, while NB consistently showed the lowest growth, suggesting media components interact with the hydrogel network to restrict bacterial growth. LB, TSB, and M1 hydrogels showed intermediate growth.

**Figure 6.**
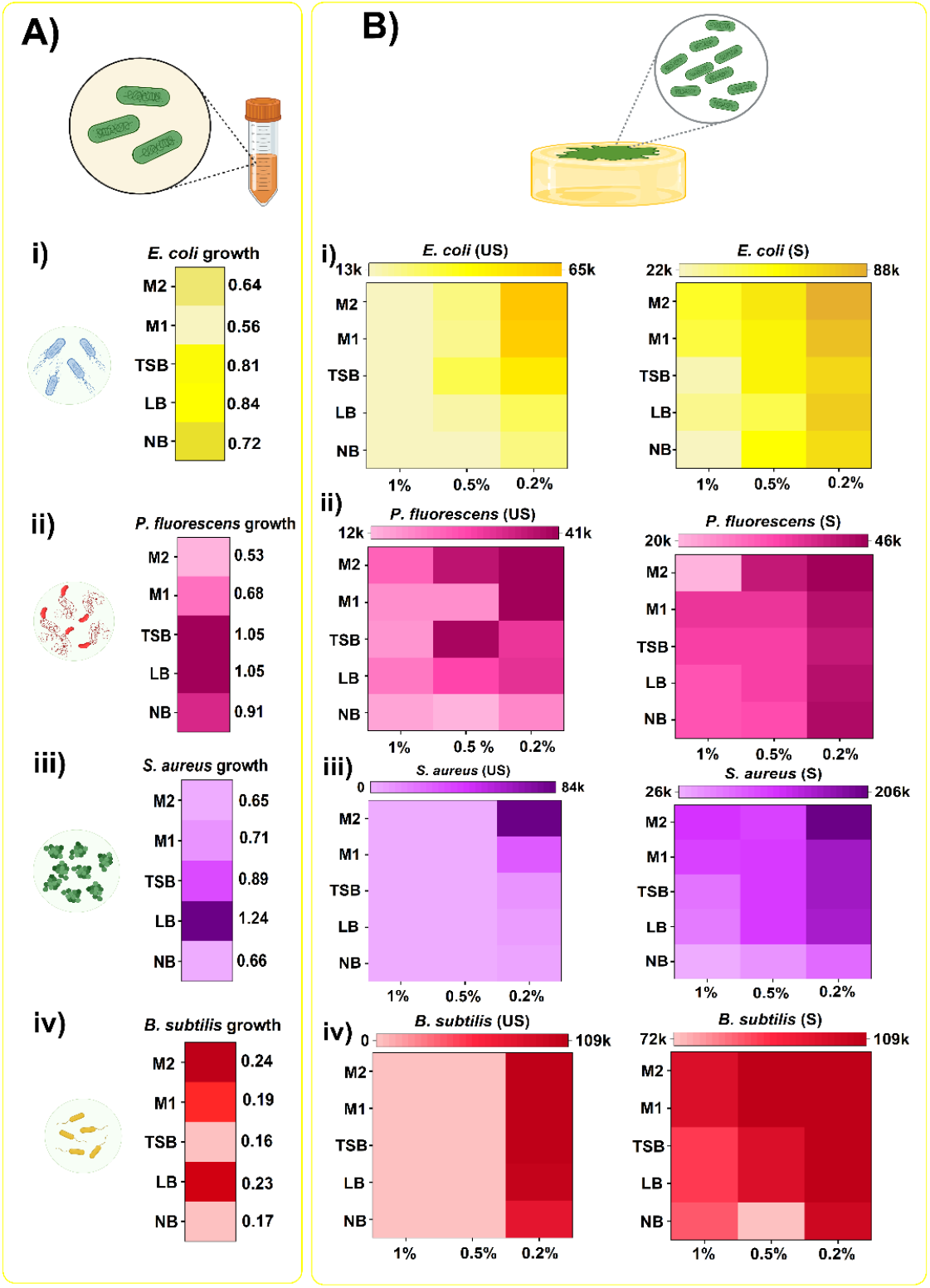
Heatmaps showing the effect of nutrient on (i) *E. coli*, (ii) *P. fluorescens*, (iii) *S. aureus*, and (iv) *B. subtilis* growth **(A)** in-solution show no consistent pattern, likely driven by media composition. (**B)** In hydrogels (1%, 0.5%, 0.2%) with encapsulated NB, LB, TSB, M1, and M2, growth is more consistent-highest in M2 across all species.

To investigate this interaction, we examined the stiffness and water loss properties of US and S agarose hydrogels encapsulated with NB, LB, TSB, M1, and M2. Compared to NB, all other media (LB, TSB, M1, and M2) resulted in a noticeable reduction in hydrogel stiffness across all concentrations (1%, 0.5%, 0.2%) with M2 producing the greatest decrease in US (Figure 7A (i-iii)) and S (Figure 7B (i-iii)). Similarly, water retention was higher (i.e., lower water loss) in hydrogels containing LB, TSB, M1, and M2, relative to NB in US (Figure 7C (i-iii)) and S (Figure 7D (i-iii)).

**Figure 7.**
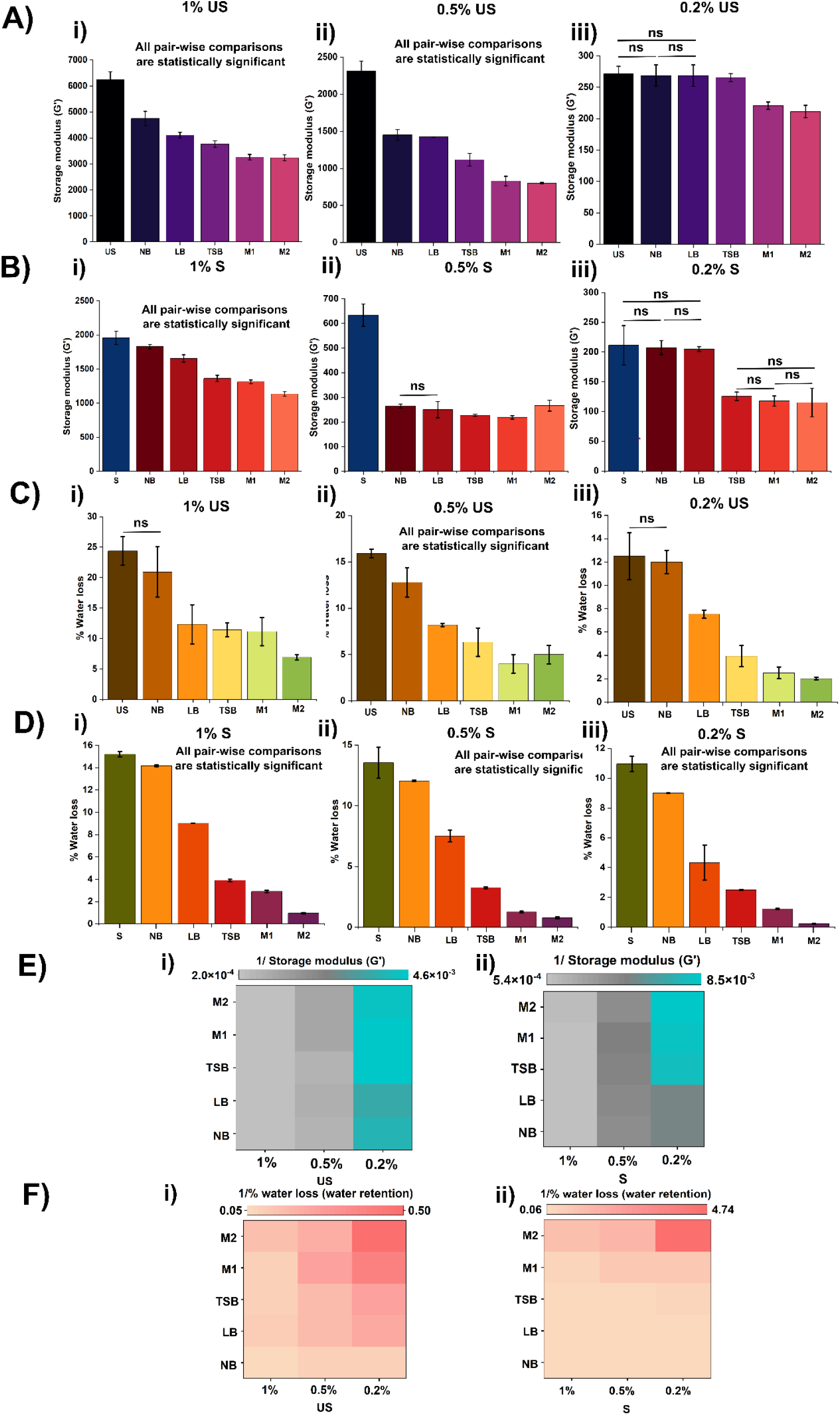
Effect of nutrient on hydrogel properties and bacterial growth **(A), (B)** Stiffness of US (i) 1%, (ii) 0.5%, and (iii) 0.2% and 1%, (ii) 0.5%, and (iii) 0.2% S hydrogels encapsulated with NB, LB, TSB, M1, and M2. **(C), (D)** % water loss of US (i) 1%, (ii) 0.5%, and (iii) 0.2% and 1%, (ii) 0.5%, and (iii) 0.2% S hydrogels encapsulated with NB, LB, TSB, M1, and M2. Data are presented as mean ± standard deviation (SD) from n=3 independent replicates. ANOVA test was performed with statistical significance set to a *P* < 0.05. Non-significant comparisons are indicated as ‘ns’. **(E)** Heatmap of elasticity (1/stiffness) and **(F)** water retention (1/water loss) shows M2-encapsulated hydrogels have highest elasticity and water retention.

Our findings indicate that encapsulated nutrients influence hydrogel stiffness and water loss, thereby affecting bacterial growth. To illustrate this, we plotted inverted stiffness (Fig.7 E (i) and (ii)) and water loss values (Fig.7 F (iii) and (iv)) as proxies for elasticity and water retention, respectively. These heatmaps reveal that media such as TSB, M1, and M2 produce more elastic and water retaining hydrogels. These trends align with conditions that support greater bacterial growth, suggesting that increased hydrogel elasticity and water retention promote bacterial proliferation.

Furthermore, we assessed the hydrophilicity of US and S agarose hydrogels. Contact angle measurements revealed that unsubstituted hydrogels (1%, 0.5%, 0.2%) exhibit moderate wettability, with contact angles of 76.6°, 62.1°, and 58.5°, respectively (Supp. Mat. Figure S25), indicating they are neither strongly hydrophobic nor fully hydrophilic. Interestingly, when encapsulated with bacterial nutrient media, these hydrogels became more hydrophilic (Supp. Movie 1-5), suggesting that encapsulation alters their surface properties. In contrast, substituted hydrogels were inherently hydrophilic, and encapsulating media did not affect their hydrophilicity (Supp. Movie 6-10).

These findings demonstrate that encapsulated nutrient media alter hydrogel stiffness, water retention, and surface wettability, thereby influencing bacterial growth.

### Unsubstituted agarose hydrogels selectively promote/inhibit bacterial growth

Interestingly, growth assays revealed that 0.5% and 1% US hydrogels promoted *E. coli* and *P. fluorescens* growth, but inhibited *S. aureus* and *B. subtilis*, while 0.2% hydrogels supported minimal growth (Figure 2, and Figure 5G and 5H). To understand this selective inhibition, we investigated bacterial cell surface properties, particularly the surface charge, and how this impacts interaction with the hydrogel matrix.

Zeta potential measurements showed that 0.5% and 1% US hydrogels are negatively charged (−1.42 ± 0.39 and −1.54 ± 0.25), while 0.2% unsubstituted and all substituted hydrogels are positively charged (Figure 8A). Cytochrome C is a small positively charged heme protein commonly used to assess surface charge; its binding to bacterial cells serves as an indicator of cell surface negativity. Cytochrome C binding assays revealed that *S. aureus* and *B. subtilis* possess highly negative surface charges (binding: 56% and 62.5%), in contrast to *E. coli* and *P. fluorescens* (43% and 32%) (Figure 8B). These findings suggest that electrostatic repulsion between negatively charged Gram-positive bacteria and negatively charged hydrogels inhibits growth. To test this hypothesis, we embedded positively charged cytochrome C into 1% unsubstituted hydrogels. This reversed the inhibitory effect and restored *B. subtilis* growth (Figure 8C), confirming that electrostatic interactions play a key role in modulating bacterial viability within hydrogels.

**Figure 8.**
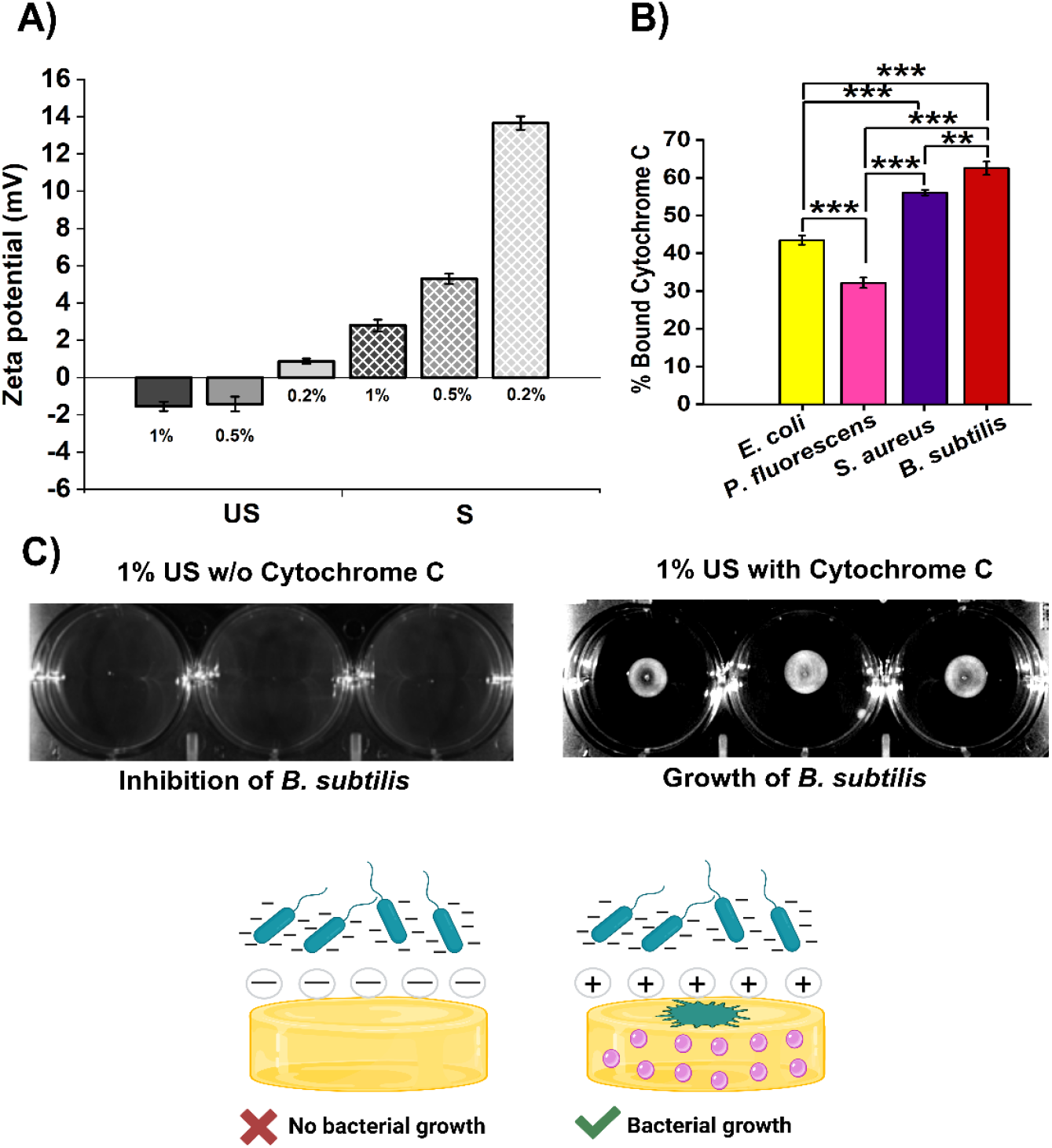
Evaluation of electrostatic interactions between bacteria and hydrogels **A)** Zeta potentials of US and S hydrogels (1%, 0.5%, and 0.2%) show surface charge differences. **B)** Cytochrome C assay reveals bacterial binding affinities. Data are presented as mean ± standard deviation (SD) from n=3 independent replicates. Statistical significance was assessed using ANOVA. P value: *P < 0.05, **P < 0.01, ***P < 0.001. **C)** *B. subtilis* growth inhibition on negatively charged 1% US hydrogels. Growth restored in 1% US hydrogel encapsulated with cytochrome C (in pink).

To evaluate whether other bacterial cell surface properties contributed to the observed selectivity, we also assessed bacterial surface hydrophobicity, membrane rigidity, and polarization (Figure S27). While minor differences were observed between species, none showed a consistent correlation with the differential growth patterns. These results further support the conclusion that electrostatic interactions are the dominant factor governing bacterial compatibility with agarose hydrogels.

## Discussion

Our study demonstrates that the physicochemical properties of agarose hydrogels play a significant role in regulating bacterial growth. By systematically varying the hydrogel type, hydrogel concentration, and the composition of the encapsulated nutrient media, we show that bacterial growth is influenced not only by nutrient availability but also by the mechanical stiffness, hydration capacity, and electrostatic characteristics of the hydrogel. These findings reveal that material properties intrinsic to agarose hydrogels can modulate bacterial growth behaviour across diverse species, highlighting the importance of considering the hydrogel matrix and surface microenvironment when designing smart bacterial or antibacterial substrates.

We show that hydrogel stiffness and water content are critical determinants of bacterial growth, with softer and more hydrated networks consistently supporting greater growth across all tested bacterial species. In our experimental setup, bacterial inoculation was performed by pricking the hydrogel surface, thereby providing entry points through which bacteria could infiltrate the matrix and establish growth both at the surface and within the hydrogel interior. This configuration bridges surface-associated and partially invasive environments and may better reflect natural settings where bacteria interact dynamically with soft materials ^16, 25, 34^. The enhanced growth in softer hydrogels suggests that reduced mechanical resistance facilitates both nutrient diffusion and spatial expansion, while higher water retention maintains a stable microenvironment that protects against desiccation. Notably, previous studies have reported inconsistent relationship between hydrogel stiffness and bacterial growth, likely due to differences in bacterial localization (surface vs. bulk matrix), unaccounted water loss characteristics, or varying definitions of mechanical compliance^26,29,31^. Together, our findings provide a mechanistic explanation for how hydrogel properties modulate bacterial growth.

To rigorously validate these findings, we performed a Type-I four-way factorial ANOVA with bacterial species, hydrogel type, polymer concentration, and encapsulated nutrient medium as independent variables. While mechanical stiffness and water retention were not included explicitly as independent factors, these physical properties are emergent from the combination of the tested variables. The statistical analysis revealed that all main effects and interaction terms were highly significant (Table S26), with the most pronounced effects arising from the species–hydrogel type interaction (F = 617.53) and a strong three-way interaction involving hydrogel concentration (F = 123.31). These results underscore the multifactorial nature of bacterial growth regulation and the importance of species-specific responses to the mechanical and hydration properties of the hydrogel matrix.

In addition to mechanical and hydration effects, our findings reveal the active and dual role of the encapsulated nutrient media, not only as a metabolic substrate for bacterial growth but also as a modulator of hydrogel physicochemical characteristics. While previous studies have shown that nutrient components like tryptone can modulate hydrogel stiffness by interacting with polymer network^25^, our work extends this understanding by demonstrating that media composition also governs the hydrogel’s water retention capacity, a factor not explicitly addressed by earlier reports. By showing that media composition influences both hydrogel stiffness and water content, we reveal how these physicochemical changes indirectly dictate bacterial proliferation. These results highlight the importance of considering medium-matrix interactions as integral to experimental design, particularly in hydrogel-based infection or colonization models.

Notably, we identify electrostatic interactions as a species-selective regulatory mechanism. Growth inhibition of Gram-positive bacteria in negatively charged unsubstituted agarose is attributable to surface charge repulsion, which was reversed by modifying the hydrogel charge via cytochrome C encapsulation. This observation highlights an underexplored dimension of hydrogel-bacteria interactions, in which physicochemical mismatch between hydrogel charge and bacterial envelope can serve as a selective growth-modulating mechanism.

Collectively, our results establish a comprehensive framework for understanding how the mechanical, hydration, and electrostatic properties of agarose hydrogels regulate bacterial growth. These findings have broad implications for the rational design of smart biomaterials that either promote or inhibit bacterial colonization. Such insights are directly relevant to the development of more physiologically relevant *in vitro* infection models, antimicrobial coatings, and biosensing platforms. Importantly, our study highlights the necessity of integrating material properties with microbiological design to harness the full potential of hydrogel-based technologies.

## Materials and Methods

### Bacterial culture

*P. fluorescens* and *S. aureus USA 300 JE2* (BEI Resources) was obtained from Warwick Medical School, and *B. subtilis 168* was obtained from Nanosyrinx, University of Warwick. The strain of *E. coli* used was Uropathogenic *E. coli* CFT073. Bacteria were revived from their glycerol stocks and streaked on lysogeny agar for *E. coli, P. fluorescens*, and *B. subtilis* and tryptic soy agar (TSA) for *S. aureus*. Single colonies were picked from each plate and inoculated into the Luria broth medium for *E. coli, P. fluorescens*, and *B. subtilis* and tryptic soy broth for *S. aureus* for primary culture at 37°C and 180 RPM overnight. The optical density (OD) of bacterial cultures was determined at 600 nm. The bacterial cultures were adjusted to an OD of 0.01 using culture medium appropriate for each bacterial species. 2 µL of these cultures were utilized for evaluating bacterial growth in agarose hydrogels.

### Synthesis of bacterial culture media encapsulated agarose hydrogels

Unsubstituted agarose hydrogels were prepared using high-melting agarose of molecular grade (BIO-41025, lot number: ES520-B103080) purchased from Meridian Biosciences, Bioline. High-melting agarose has a melting temperature of 88.8°C and gelling temperature of 38.5°C. Substituted agarose hydrogels were prepared using low-melting agarose of hydroxyethyl type (product number: A9414, CAS number: 39346-81-1) purchased from Sigma-Aldrich, Merck. Low-melting agarose has a melting temperature of ~65°C and gelling temperature of 26-30°C.

Agarose hydrogels were prepared in batches of 50 mL for each type of nutrient media and concentrations: 0.5 g agarose in 50mL (1%), 0.25 g agarose in 50 mL (0.5%), 0.1 g in 50 mL (0.2%) for NB, LB, TSB, M1, and M2 media. The solutions were heated up to 100°C until all the agarose was completely dissolved (~ < 2 minutes). 2mL of these agarose solutions were pipetted onto sterile tissue culture graded 6-well plates (Greiner) and allowed to gel at room temperature. For each of these conditions, the experiments were performed in three replicates. To ensure complete hydrogel solidification, the plates were used for experiments after 45 minutes of gelation.

### Bacterial growth in hydrogels

Once the hydrogels were solidified, 2 µL of bacterial cultures were introduced separately for each condition. The bacterial culture was pricked at the centre of the agarose hydrogel in 6-well plate such that the prick penetrates the gel but does not pierce the hydrogel bottom. Upon bacterial introduction on hydrogels, the plates were incubated without any disturbance of physical movements at 37°C for 18 hours. Bacterial growth was qualitatively visualized using the BIO-RAD ChemiDoc^™^ MP imaging system using the colorimetric filter with an exposure time of 50 seconds. For *B. subtilis*, the exposure time was prolonged to 1.50 minutes due to their growth characteristics, which was not apparent in 50 seconds.

### Fiji analysis

The colorimetric images were analysed in the Fiji software. All images were converted to 8-bit and a background subtraction was performed. The images were then adjusted by thresholding to focus on bacterial growth and rule out non-bacterial growth spots. After thresholding, the images were converted to the binary format for ease of segmentation and quantitative measurements of bacterial growth areas. After this, the bacterial growth areas were measured and analysed.

### Hydrogel stiffness – rheology

The stiffness of unsubstituted and substituted agarose hydrogels was determined by oscillatory rheology at 25°C by monitoring the storage modulus (Young’s modulus) (G’) and loss modulus (G’’) using the parallel plate geometry in the Anton Paar MCR 302 rheometer. First, amplitude sweeps were performed at a constant frequency of ω = 10 rad/sec to determine the linear viscoelastic regions (LVE) for each concentration and hydrogel type. Frequency sweep analysis was performed in the angular frequency range of ω = 1-100 rad/sec with a constant strain of γ = 0.05% as determined from the amplitude sweep analysis using RheoCompass software.

### Hydrogel hydrophilicity - contact angle measurements

The hydrophilicity of hydrogels was determined by contact angle measurements using the KRÜSS drop shape analyser 100. 5 µL of water was used for the measurements. KRÜSS DSA3 software with Young-Laplace mathematical model was used for recording the contact angle measurements.

### Qualitative assessment of hydrogel porosity - Cryo-SEM

Cryo-SEM was performed using a Zeiss Crossbeam 550 (Carl Zeiss, Oberkochen, Germany) Focused Ion Beam Scanning Electron Microscope (FIB-SEM) at an accelerating voltage of 2kV. The samples were maintained at cryogenic temperatures of −150°C using a Quorum 3010 cryo stage (Quorum technologies, Loughton, UK). The samples were secured into slots in aluminium stubs mounted into the cryo-shuttle and plunged into slushy nitrogen and allowed to freeze. The cryo-sludge with samples was transferred to the preparation chamber and the samples freeze-fractured by a cooled blade. The fractured surface was sublimed at −90°C for 30 minutes (etching). The temperature of the shuttle was returned to −150°C and the samples were sputter coated with platinum at 10 mA for 60 seconds. Samples were transferred to the main stage of the microscope and imaged using a range of detection modes including the Secondary Electrons Secondary Ions (SESI) and in-Lens detectors at 2kV accelerating voltage. Standard imaging magnifications were used to allow comparison of different samples.

### Water loss analysis

After hydrogel synthesis as described in 3.3.2, the 6-well plates were incubated at 37°C for 18 hours, mimicking the experimental conditions of bacterial growth on hydrogels. The weight of hydrogels before and after incubation for 18 hours were recorded. The % water loss was calculated by:

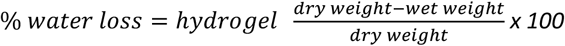

### Heatmaps and statistical analysis

All bar graphs and heatmaps were produced in OriginPro 2021b software.

A Type-I four-way factorial analysis of variance (ANOVA) was used to evaluate the effects of bacterial species, hydrogel type, polymer concentration, and encapsulated nutrient medium on bacterial growth. All main effects and interaction terms were included in the model. While hydrogel stiffness and water retention were not explicitly modelled as independent variables, they were treated as emergent properties resulting from the experimental design. ANOVA was performed using custom scripts written in R statistical programming language, which are publicly available at https://github.com/dtaylor3509/BacterialGrowthInHydrogels/tree/main.

The relationship between bacterial growth and hydrogel properties (stiffness and water loss) was assessed using Spearman’s rank correlation. The storage modulus and % water loss measured values were used for correlation with bacterial growth.

### Hydrogel charges - Zeta potential analysis

The hydrogel surface charges were determined by measuring and analyzing the zeta potential of hydrogels using the Anton Paar SurPASS 3 electrokinetic analyser. Discs with 1 cm diameter were punched out of agarose hydrogels and mounted into the recess of the cylindrical cell of the SurPass 3 instrument. The hydrogel discs were perforated with a tweezer to enable a continuous and laminar flow through the hydrogel. The discs were first put through five rinse cycles of 100 mL each at pressure of 600 – 200 mbar at a permeability index of 100-110 before zeta potential analysis. The zeta potential measurements were then determined at 25°C with 10 mM potassium chloride in water and 10 mM PBS.

### Bacterial cell membrane charges - Cytochrome C assay

Primary cultures of *E. coli, P. fluorescens, S. aureus*, and *B. subtilis* were prepared as described in 3.3.1. The cultures were pelleted at 8000 RPM for 5 minutes and the pellet was washed with 20 mM (3-(N-morpholino) propanesulfonic acid) (MOPS) buffer, pH = 7. The cells were subsequently resuspended in MOPS buffer at an OD_600nm_ = 1. 50 µg/mL cytochrome C was added to the solution and incubated for 15 minutes at room temperature. The cells were pelleted at 8000 RPM for 5 minutes and the supernatant was used for spectrophotometric detection at OD_410_ using the FLUOstar OMEGA plate reader (BMG Labtech, UK) for measuring the unbound cytochrome C.

## Supporting information

E.coli growth on 1 percent US

S. aureus growth on 1 percent US

## Author contributions

Andrea Dsouza: conceptualization, methodology, investigation, formal analysis, writing – original draft and editing; Dylan Taylor: coding, writing – review and editing, formal analysis; Chris Parmenter: methodology, investigation; Rachel A. Hand: writing - review and editing, methodology; Julia Brettschneider: formal analysis, writing – review and editing; Meera Unnikrishnan: methodology, formal analysis, writing - review and editing; Chrystala Constantinidou: funding, methodology, formal analysis, writing – review and editing; Jérôme Charmet: conceptualization, funding, methodology, formal analysis, writing – original draft, review and editing.

## Acknowledgements

We thank the Polymer Synthesis & Characterisation Research Technology Platform, The University of Warwick for their support in rheology characterization and contact angle measurements. We are also thankful to Dr Spyros Efstathiou for discussions on rheology data analysis and Dr Wai Hin Lee for help with the drop shape analyser. We are thankful to the Nanoscale and Microscale Research Centre, The University of Nottingham for access to the Cryo-SEM imaging facility. We would also like to thank Anton Paar for the loan of the SurPASS 3 instrument. Andrea would like to thank the Chancellor’s International Scholarship (2020-2024) awarded by the University of Warwick to undertake this work. Figures were created and produced in OriginPro 2021b, Biorender and Inkscape software.

