## Supplementary information for "Dynamic Bacterial Growth Modulation in Structurally Distinct and Functionally Tuneable Agarose Hydrogels"

Prof. Meera Unnikrishnan

Division of Biomedical Sciences, Warwick Medical School, The University of Warwick, Coventry, CV4 7AL, United Kingdom

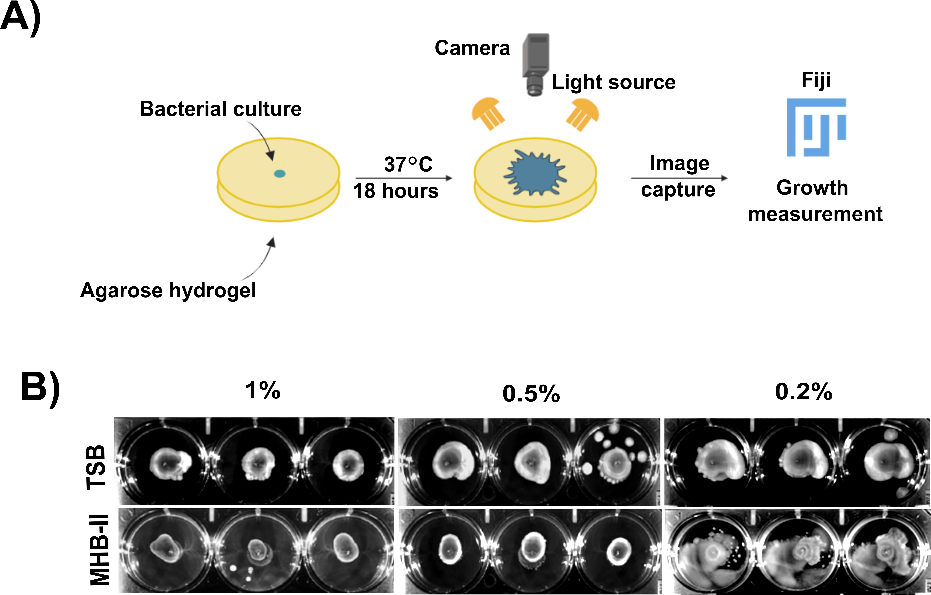

**Figure S1. A)** Schematic diagram of the experimental set-up. 3 µL of bacterial culture was introduced at the hydrogel center by pricking the surface with a pipette tip. Hydrogels were incubated at 37 °C for 18 h, imaged using a Bio-Rad gel doc system with a colorimetric filter, and bacterial growth was quantified using Fiji software. **B)** Representative images of uropathogenic *E. coli* growth in TSB and M2 encapsulated US hydrogels at 1%, 0.5% and 0.2% hydrogel concentrations.

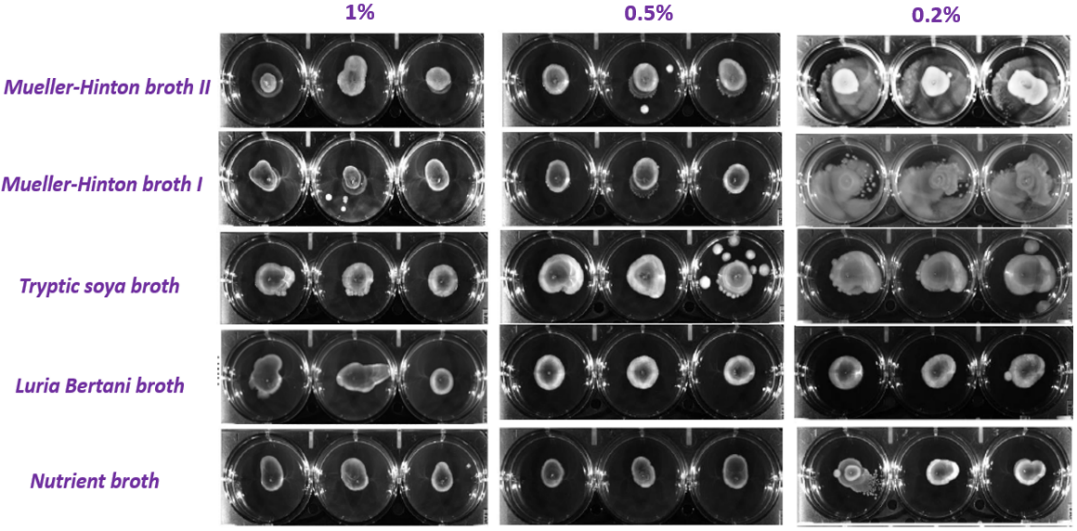

**Figure S2.** Images of uropathogenic *E. coli* growth on unsubstituted agarose hydrogels

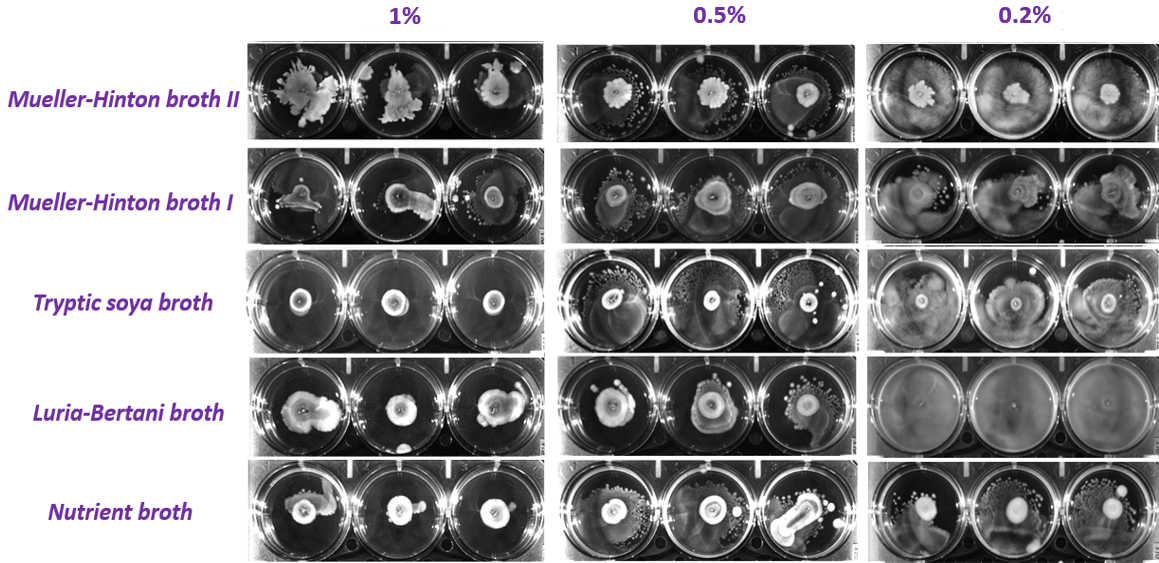

**Figure S3.** Images of uropathogenic *E. coli* growth on substituted agarose hydrogels

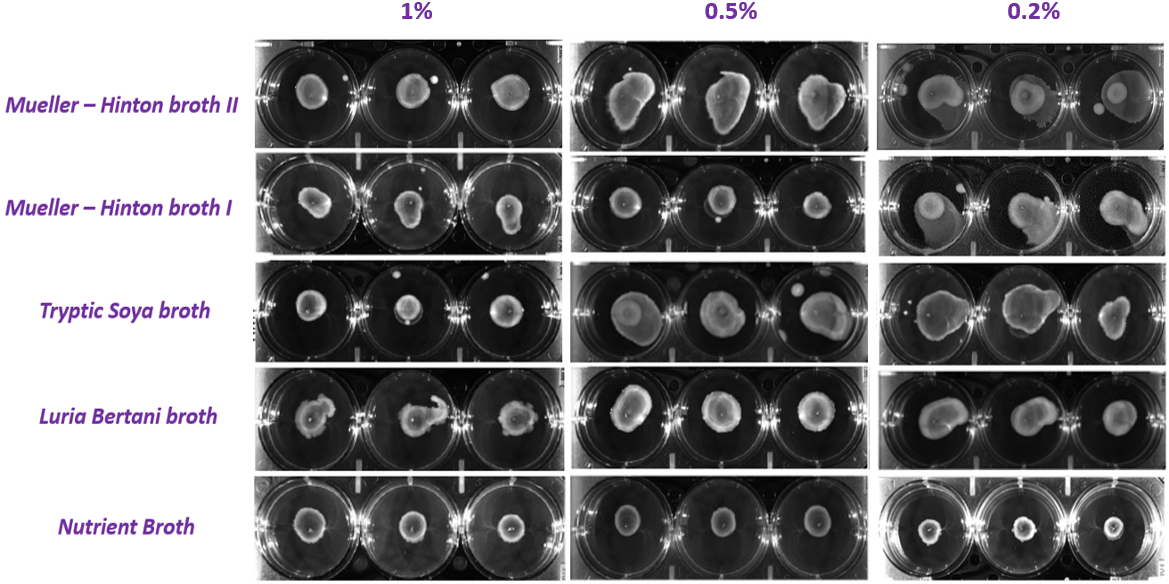

**Figure S4.** Images of uropathogenic *P. fluorescens* growth on unsubstituted agarose hydrogels encapsulated with NB, LB, TSB, M1, and M2.

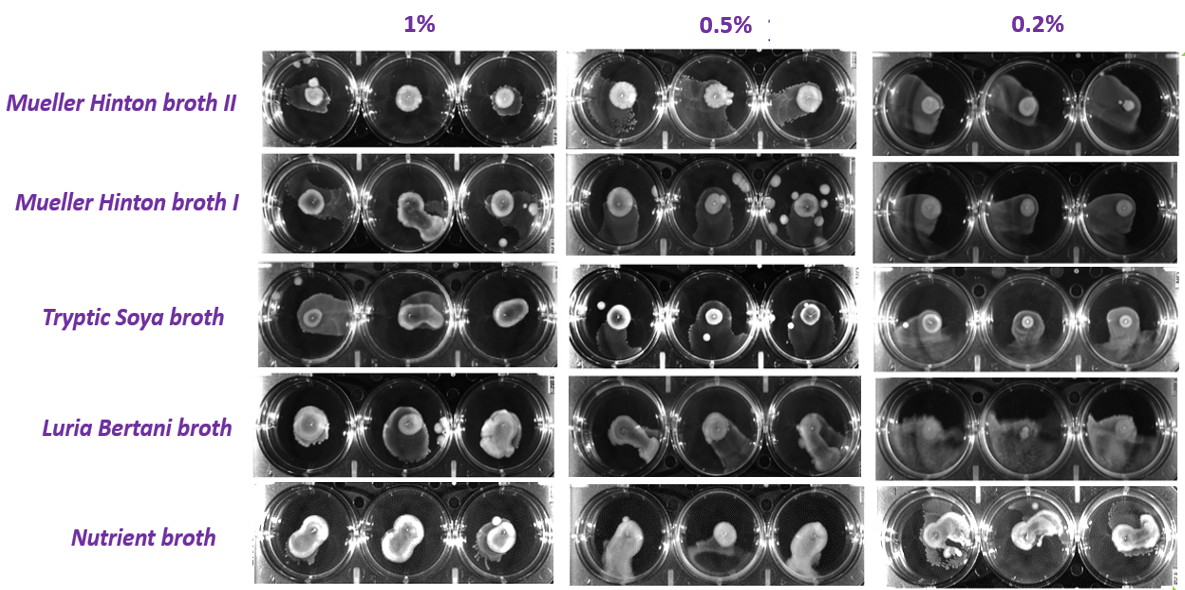

**Figure S5.** Images of uropathogenic *P. fluorescens* growth on substituted agarose hydrogels encapsulated with NB, LB, TSB, M1, and M2.

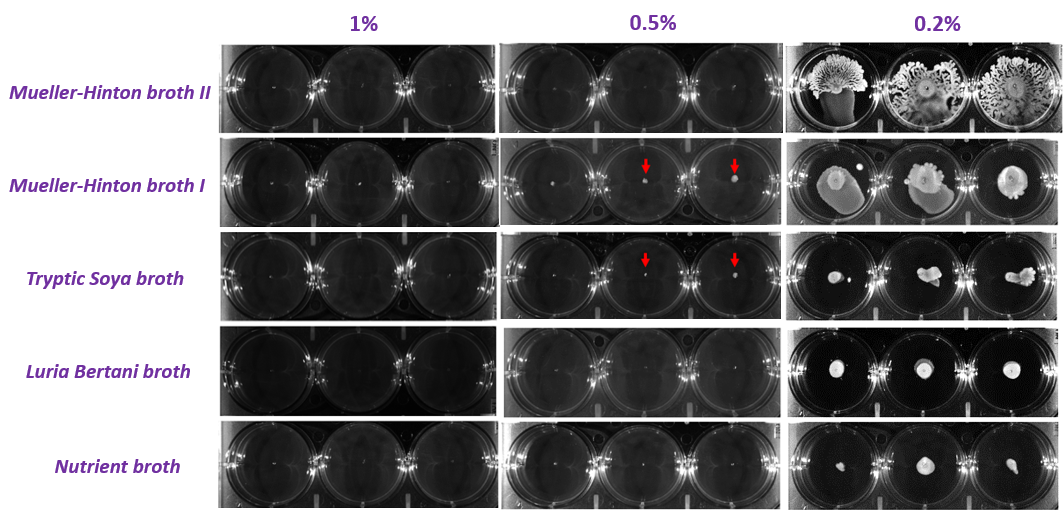

**Figure S6.** Images of uropathogenic *S. aureus* growth on unsubstituted agarose hydrogels encapsulated with NB, LB, TSB, M1, and M2. 1% and 0.5% do not support *S. aureus* growth. Red arrows indicate spotted growth at the point of inoculation.

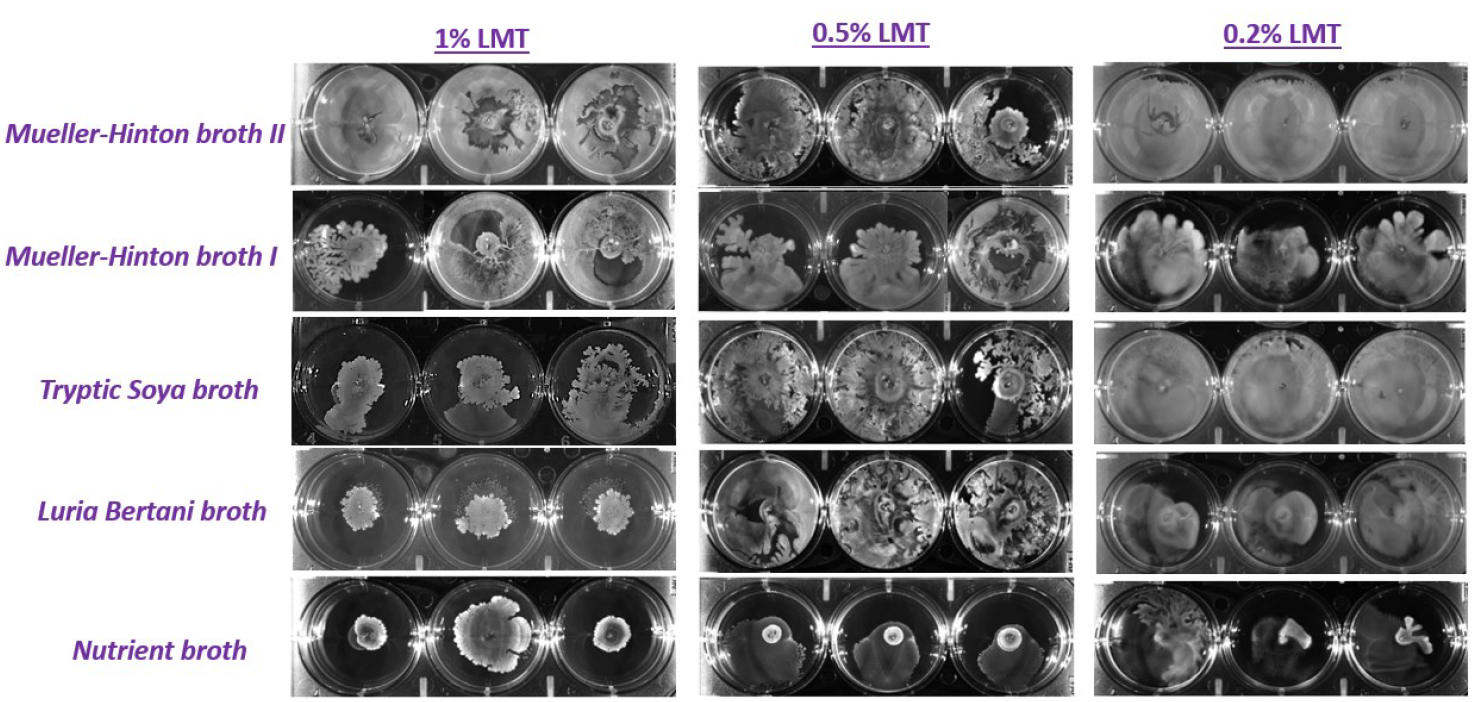

**Figure S7.** Images of uropathogenic *S. aureus* growth on substituted agarose hydrogels encapsulated with NB, LB, TSB, M1, and M2.

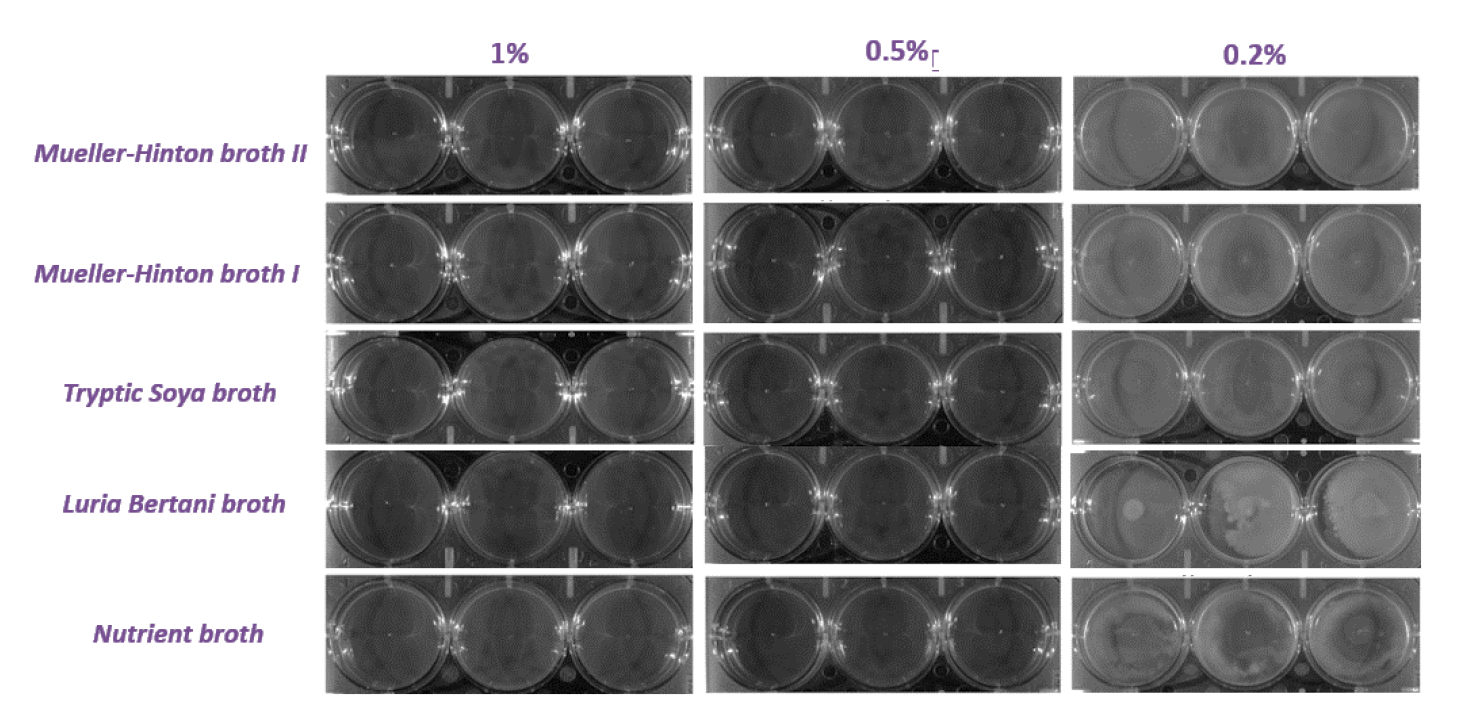

**Figure S8.** Images of uropathogenic *B. subtilis* growth on unsubstituted agarose hydrogels encapsulated with NB, LB, TSB, M1, and M2. 1% and 0.5% do not support any growth. 0.2% supports *B. subtilis* growth characterized by diffused bacterial growth in the hydrogel.

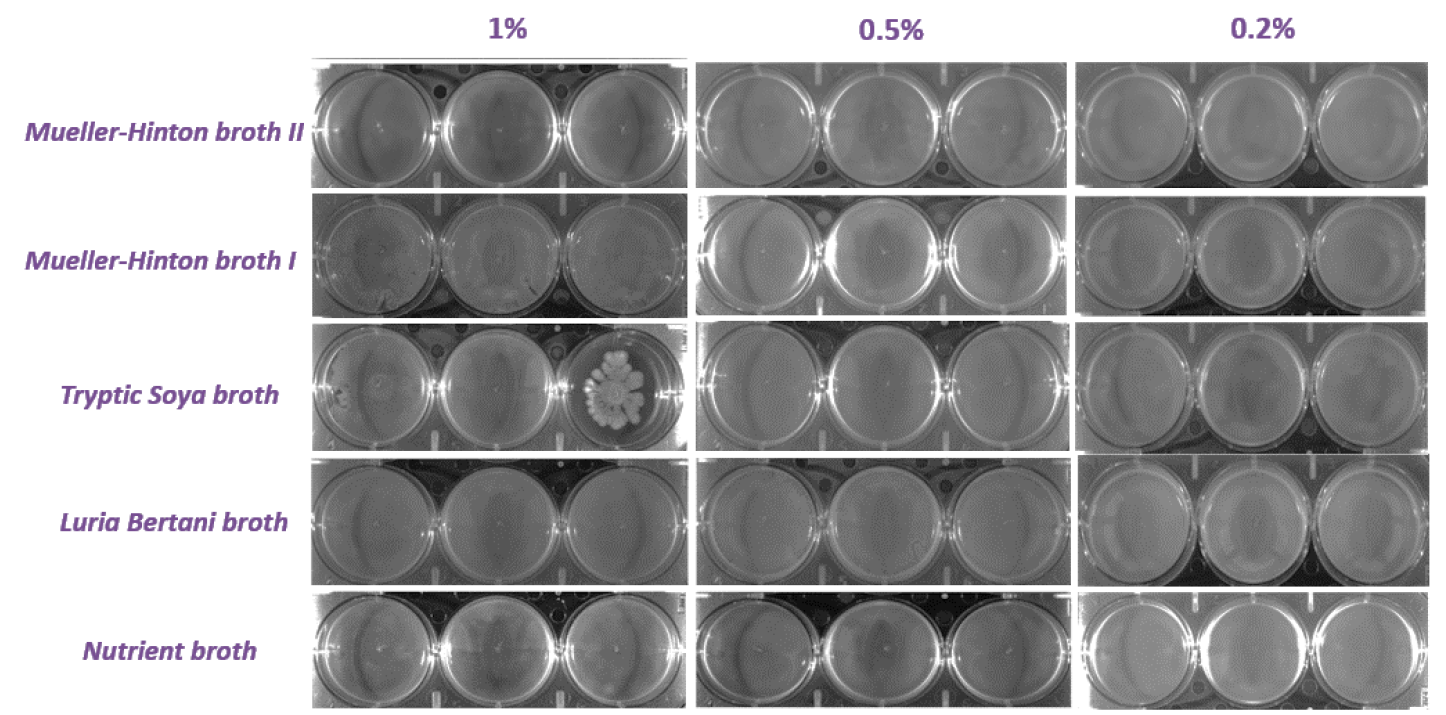

**Figure S9.** Images of uropathogenic *B. subtilis* growth on substituted agarose hydrogels encapsulated with NB, LB, TSB, M1, and M2. 1%, 0.5%, and 0.2% support *B. subtilis* growth characterized by diffused growth patterns.

| 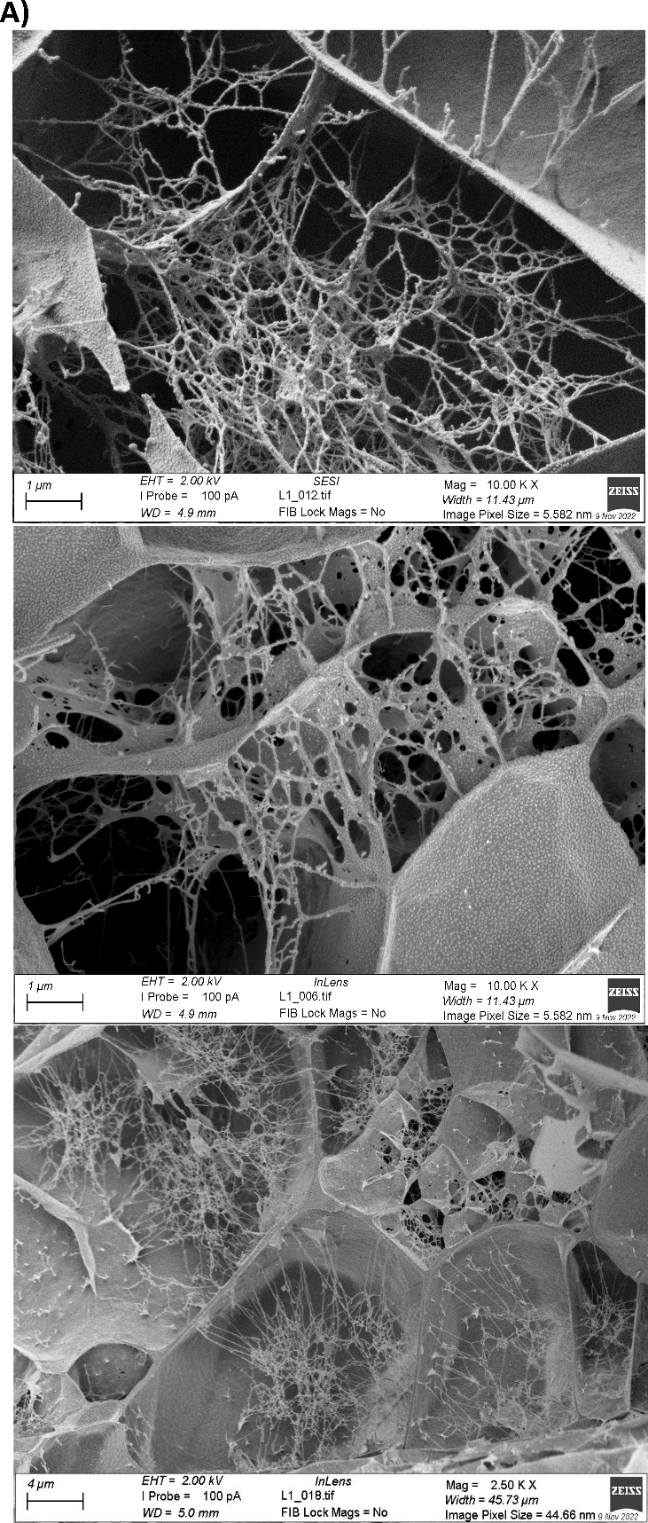 | 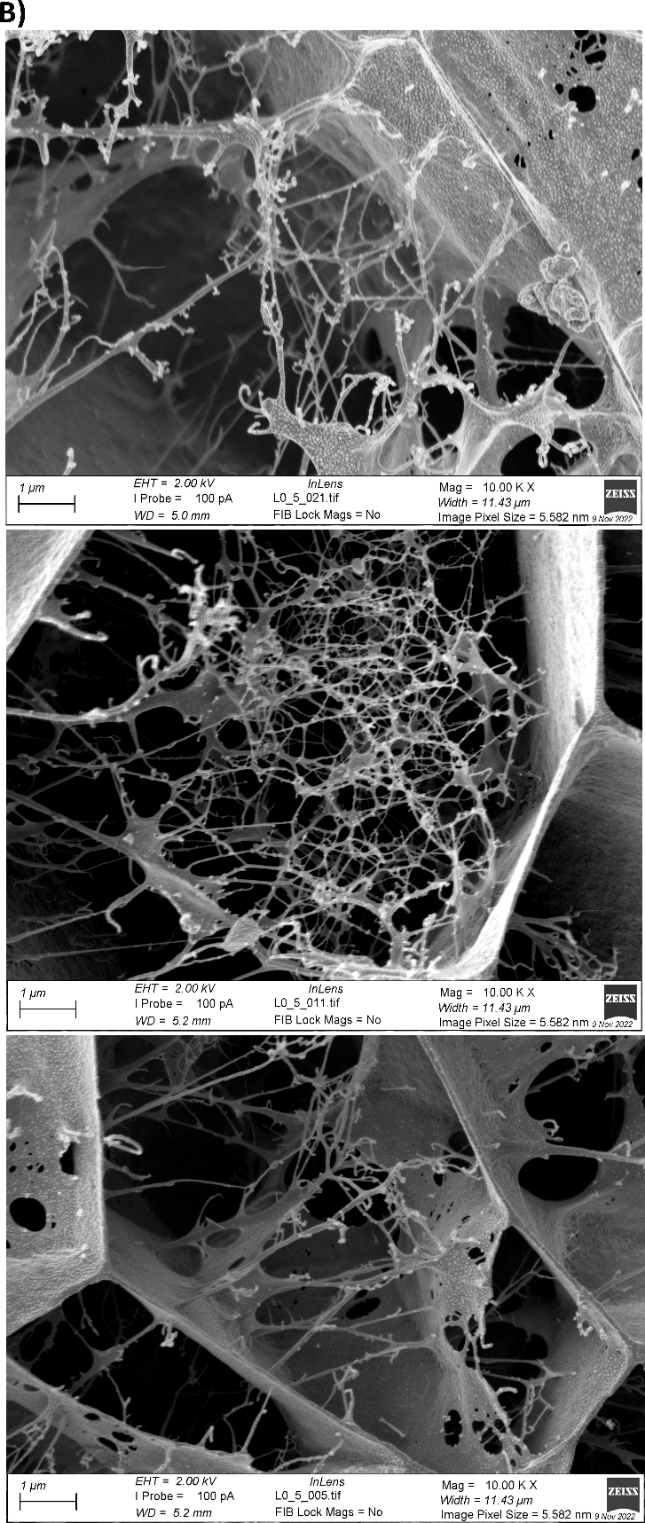 |
| --- | --- |

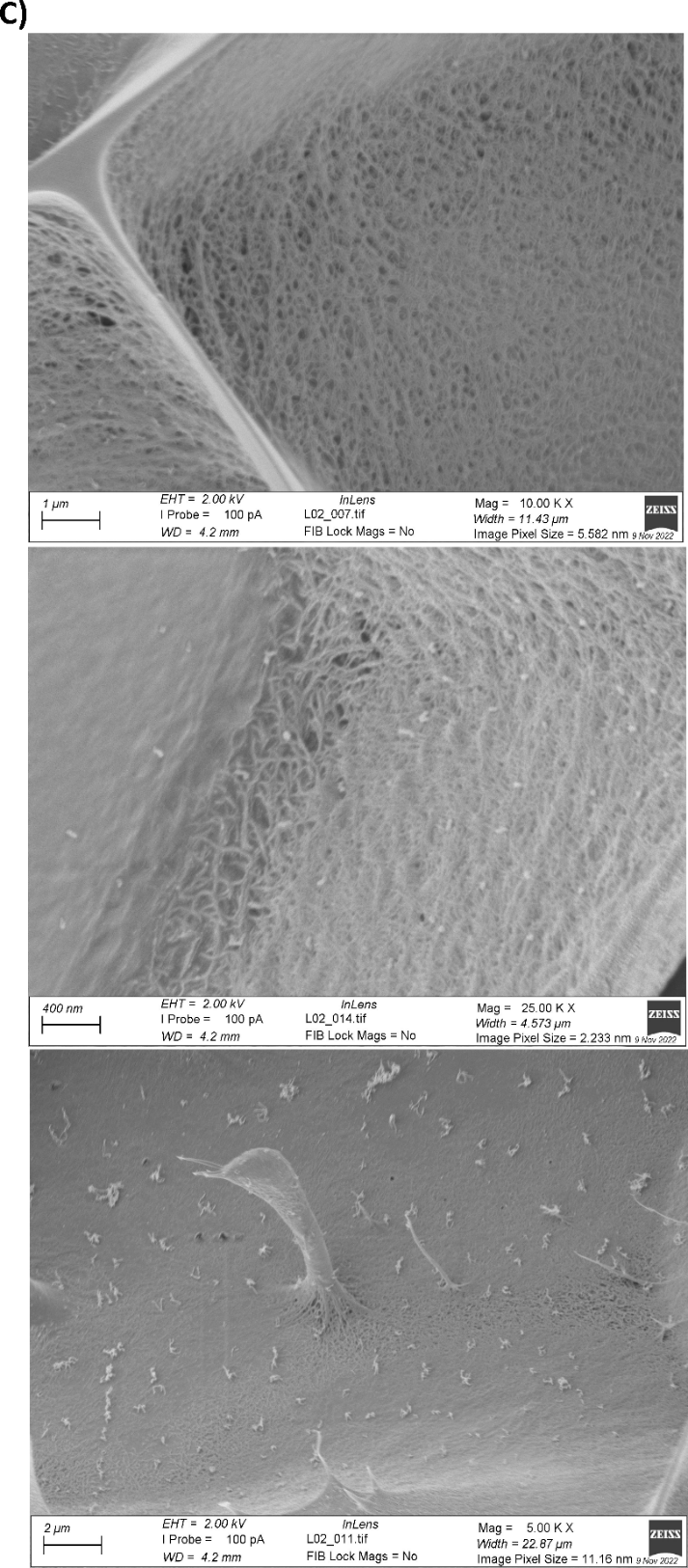

**Figure S10**. Cryo-SEM images of A) 1%, B) 0.5%, and C) 0.2% substituted agarose hydrogels. These hydrogels appear to lack pores/pore-like structures with prominent thread-like structures in all concentrations. Because 0.2% substituted hydrogels are highly hydrophilic, the samples are prone to a higher degree of ice-crystal formation. We assume that the thread-like structures have collapsed and attached onto the ice-crystals as seen in the images (C).

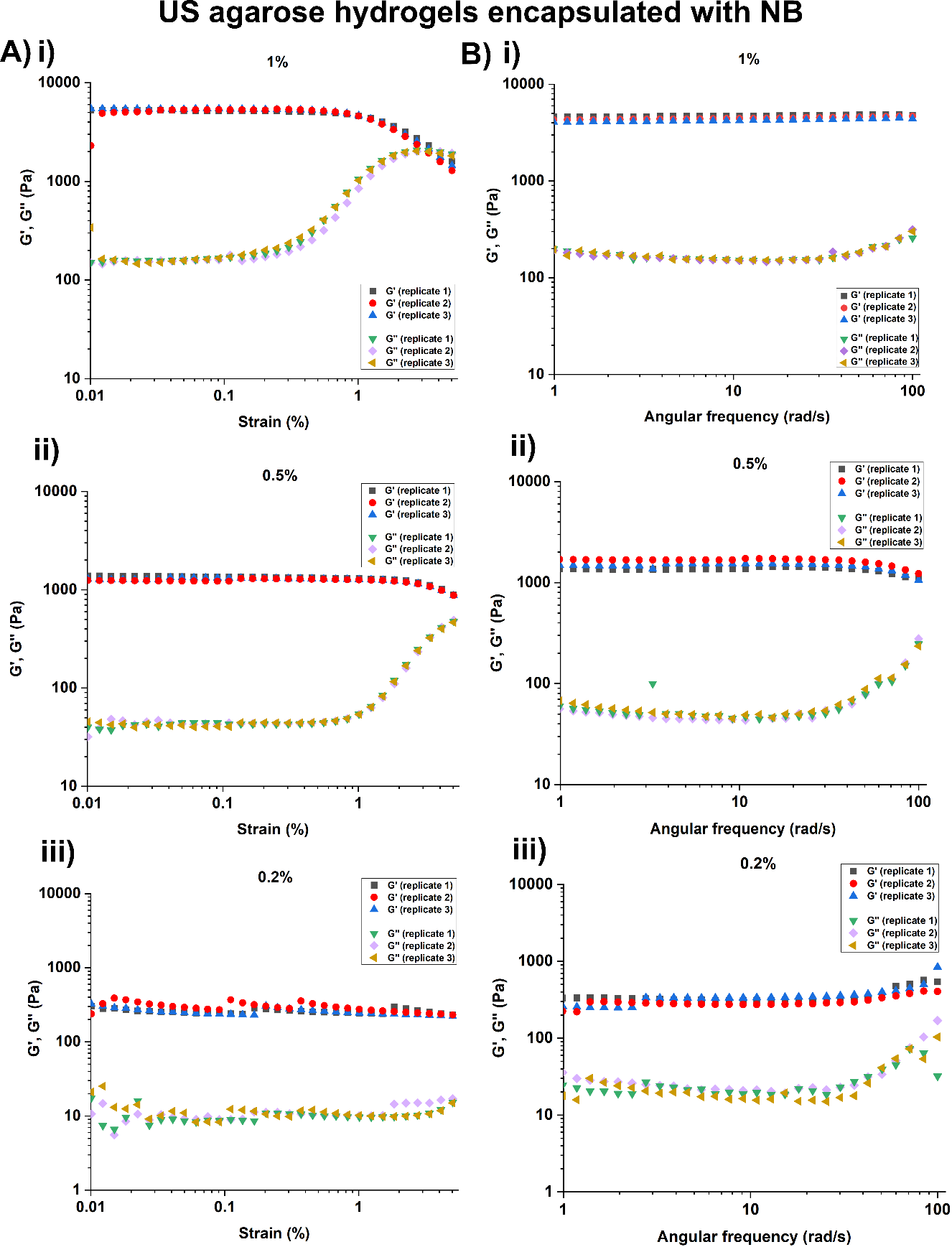

**Figure S11.** Rheology analysis. (A) Amplitude and (B) frequency sweeps of unsubstituted agarose hydrogels encapsulated with NB. All experiments were performed in three replicates – (i), (ii), (iii).

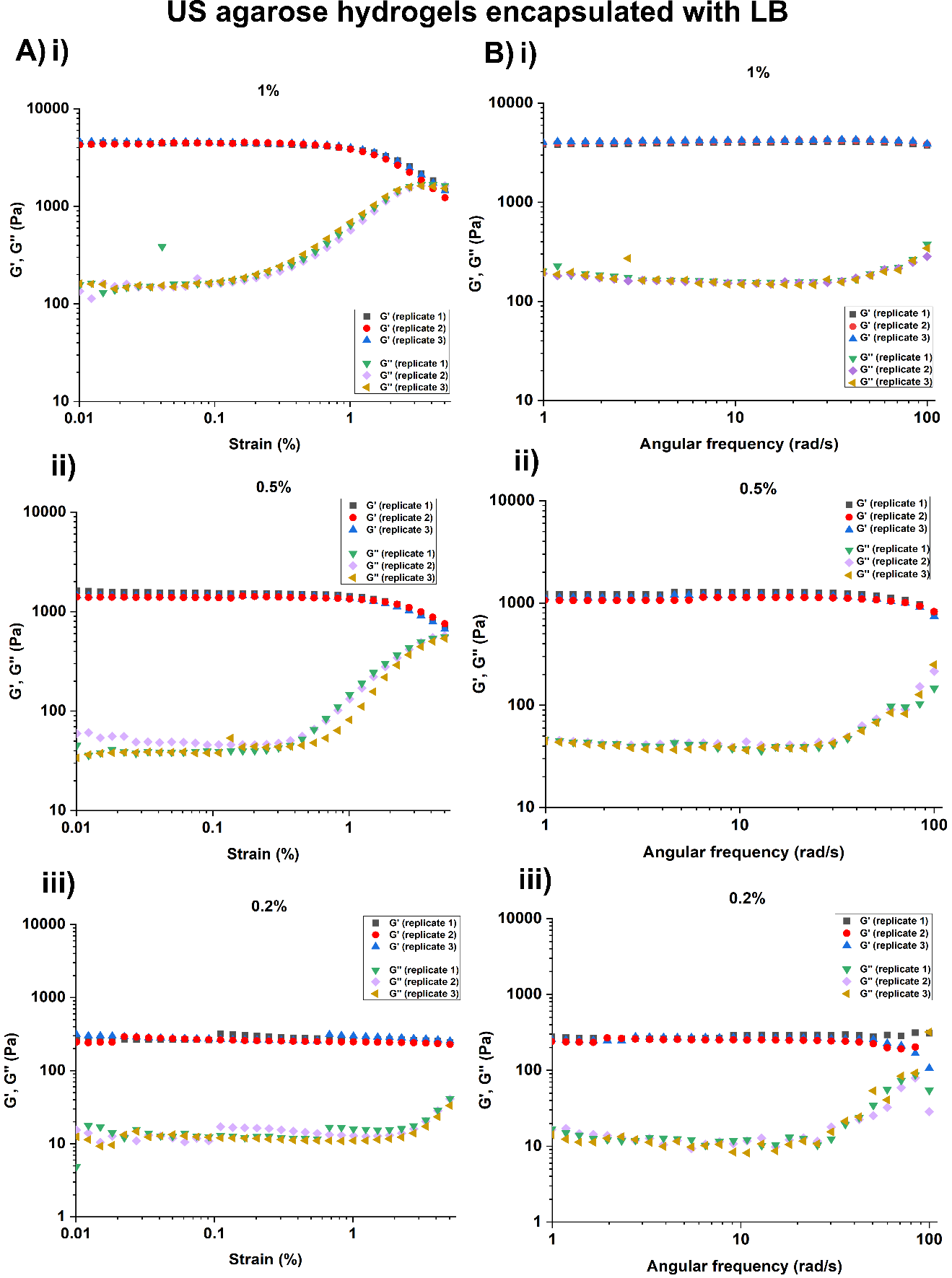

**Figure S12**. Rheology analysis. (A) Amplitude and (B) frequency sweeps of unsubstituted agarose hydrogels encapsulated with LB. All experiments were performed in three replicates – (i), (ii), (iii).

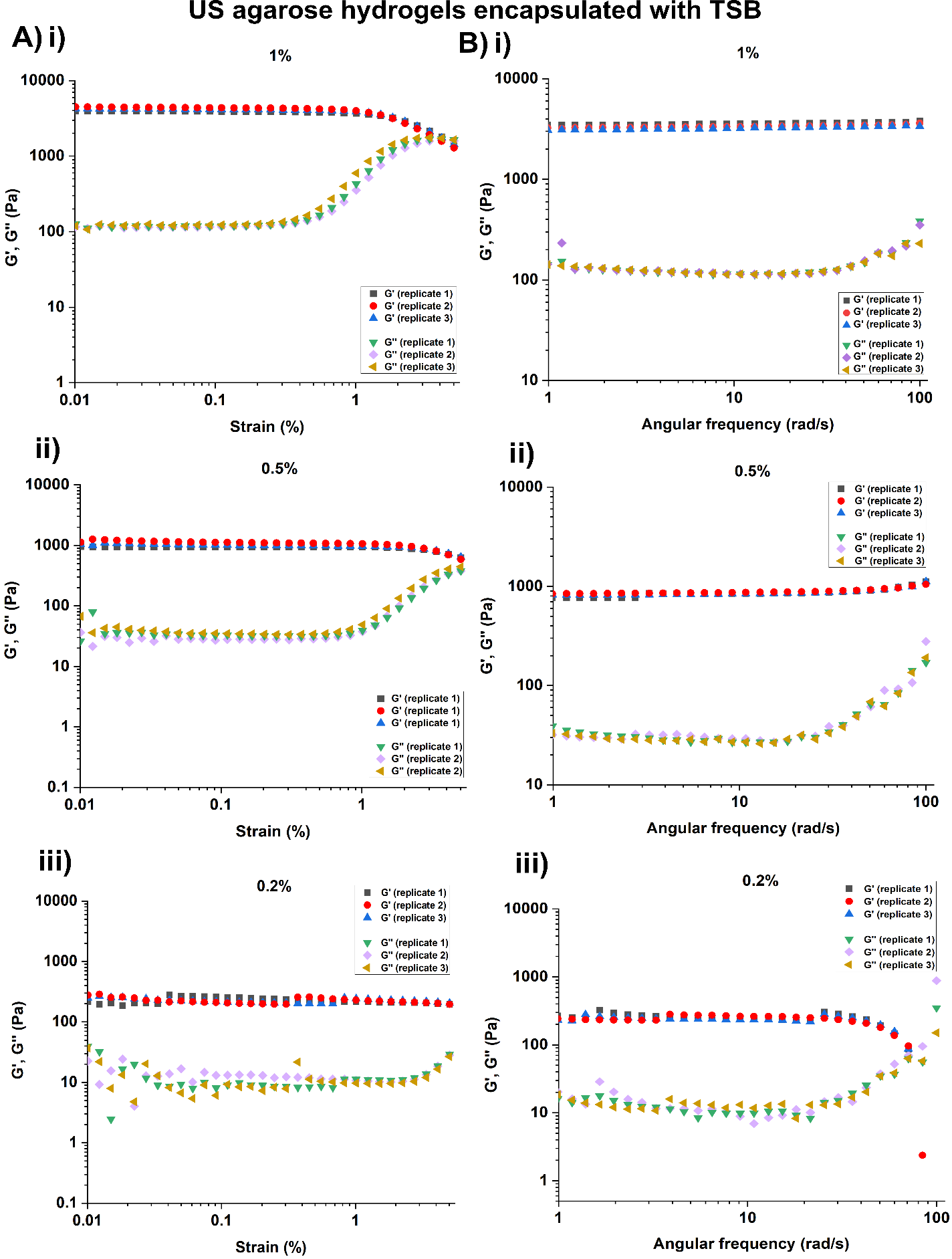

**Figure S13.** Rheology analysis. (A) Amplitude and (B) frequency sweeps of unsubstituted agarose hydrogels encapsulated with TSB. All experiments were performed in three replicates – (i), (ii), (iii).

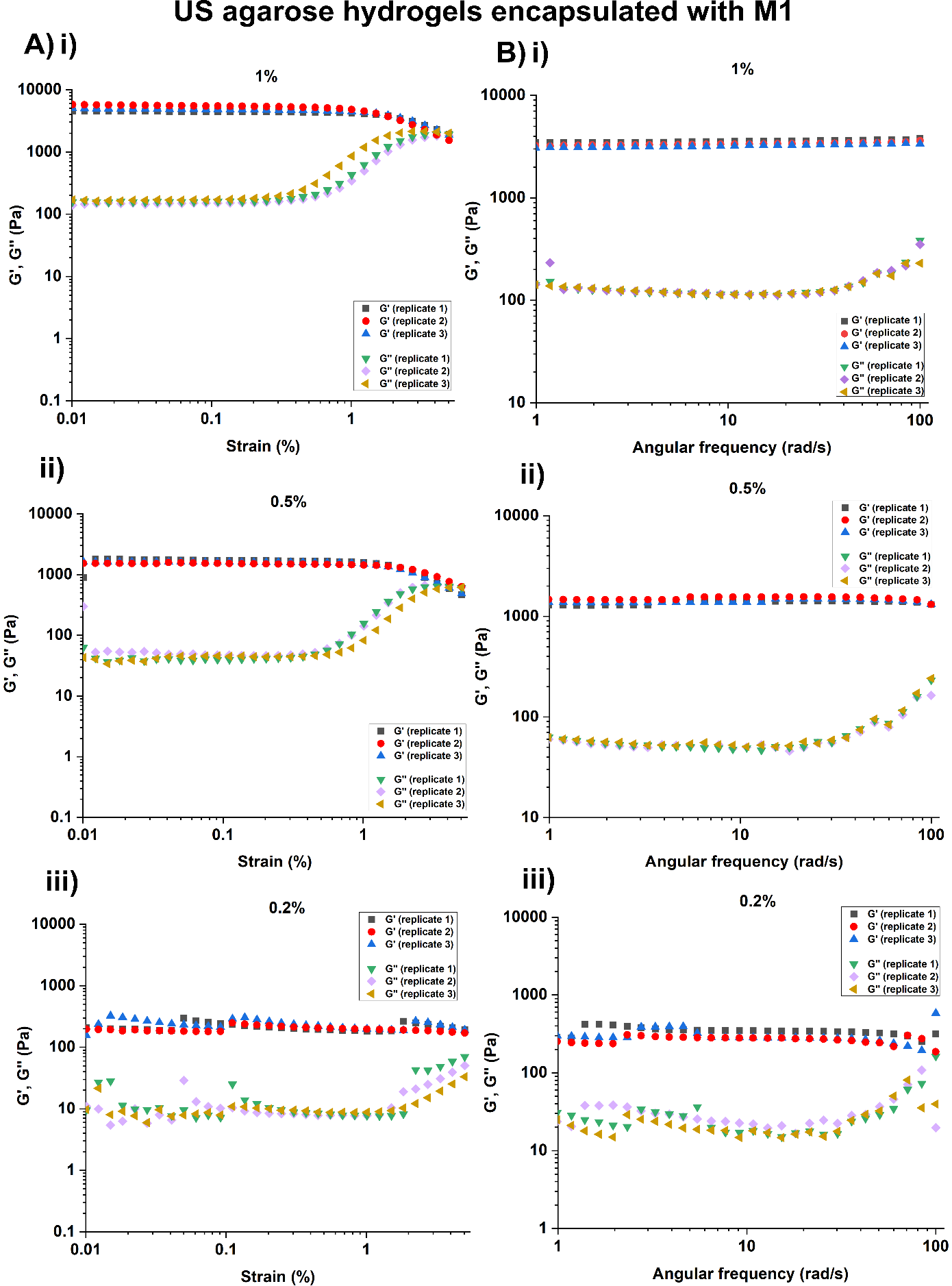

**Figure S14.** Rheology analysis. (A) Amplitude and (B) frequency sweeps of unsubstituted agarose hydrogels encapsulated with M1. All experiments were performed in three replicates – (i), (ii), (iii).

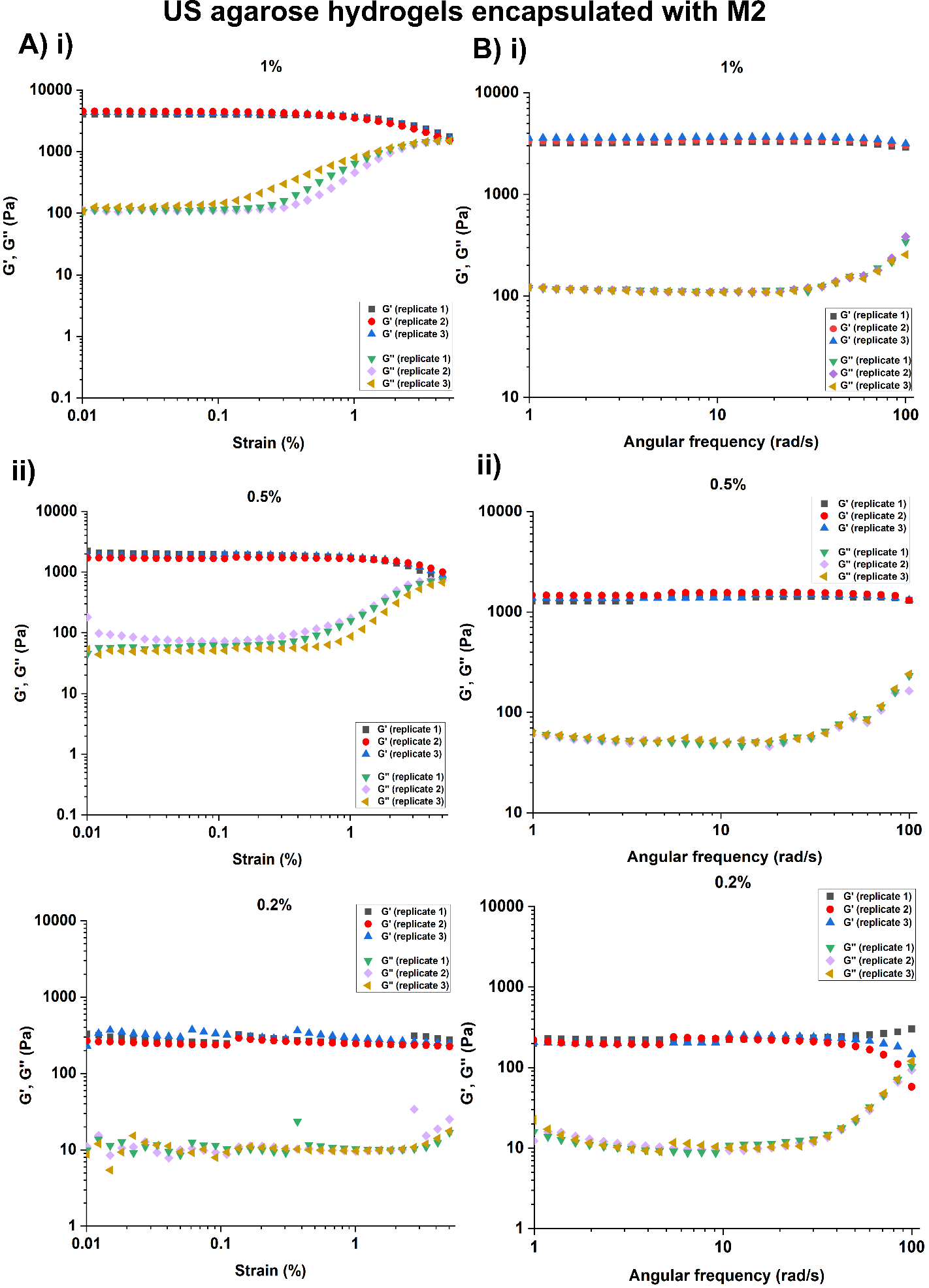

**Figure S15.** Rheology analysis. (A) Amplitude and (B) frequency sweeps of unsubstituted agarose hydrogels encapsulated with M2. All experiments were performed in three replicates – (i), (ii), (iii).

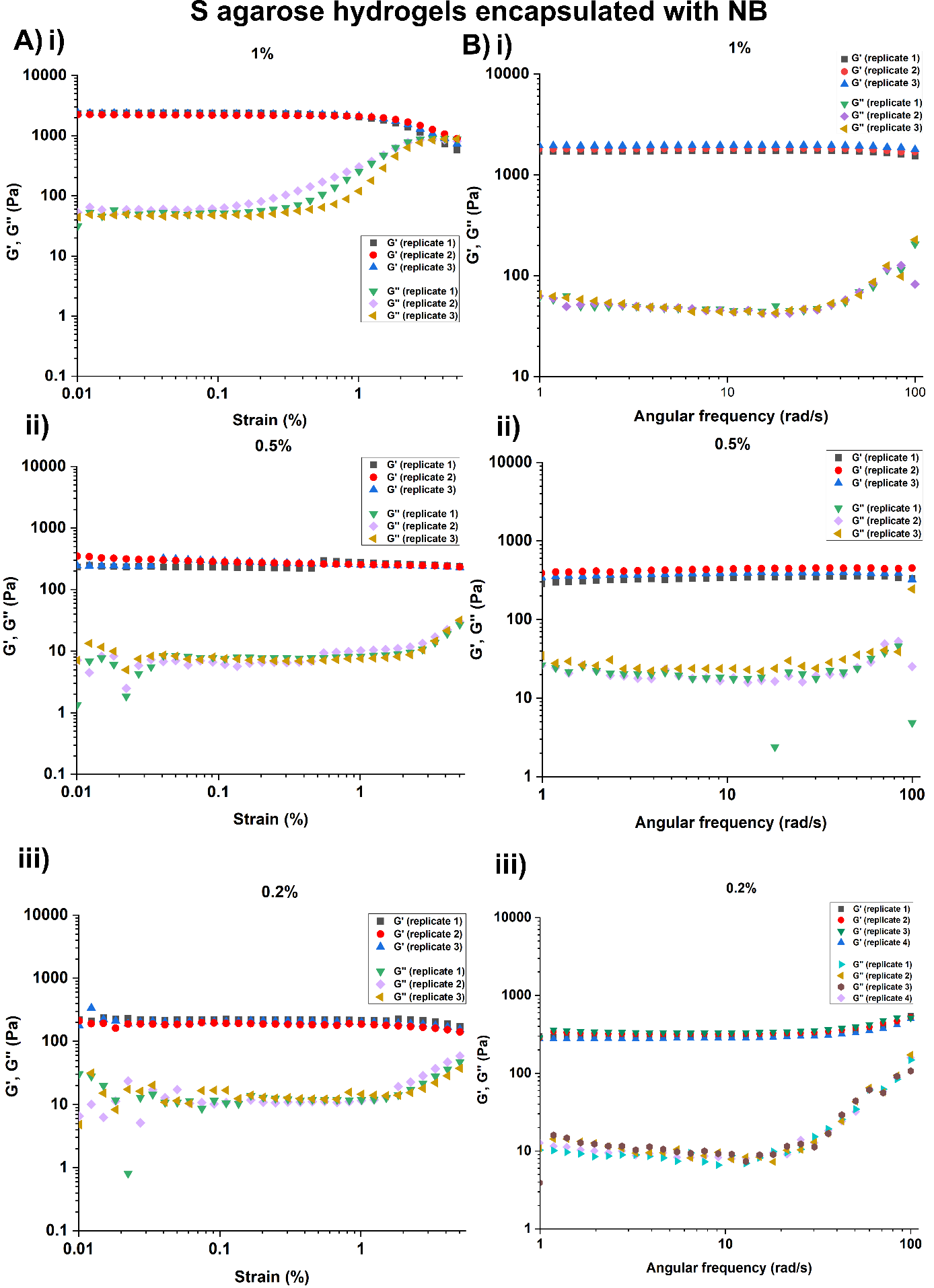

**Figure S16.** Rheology analysis. (A) Amplitude and (B) frequency sweeps of substituted agarose hydrogels encapsulated with NB. All experiments were performed in three replicates – (i), (ii), (iii).

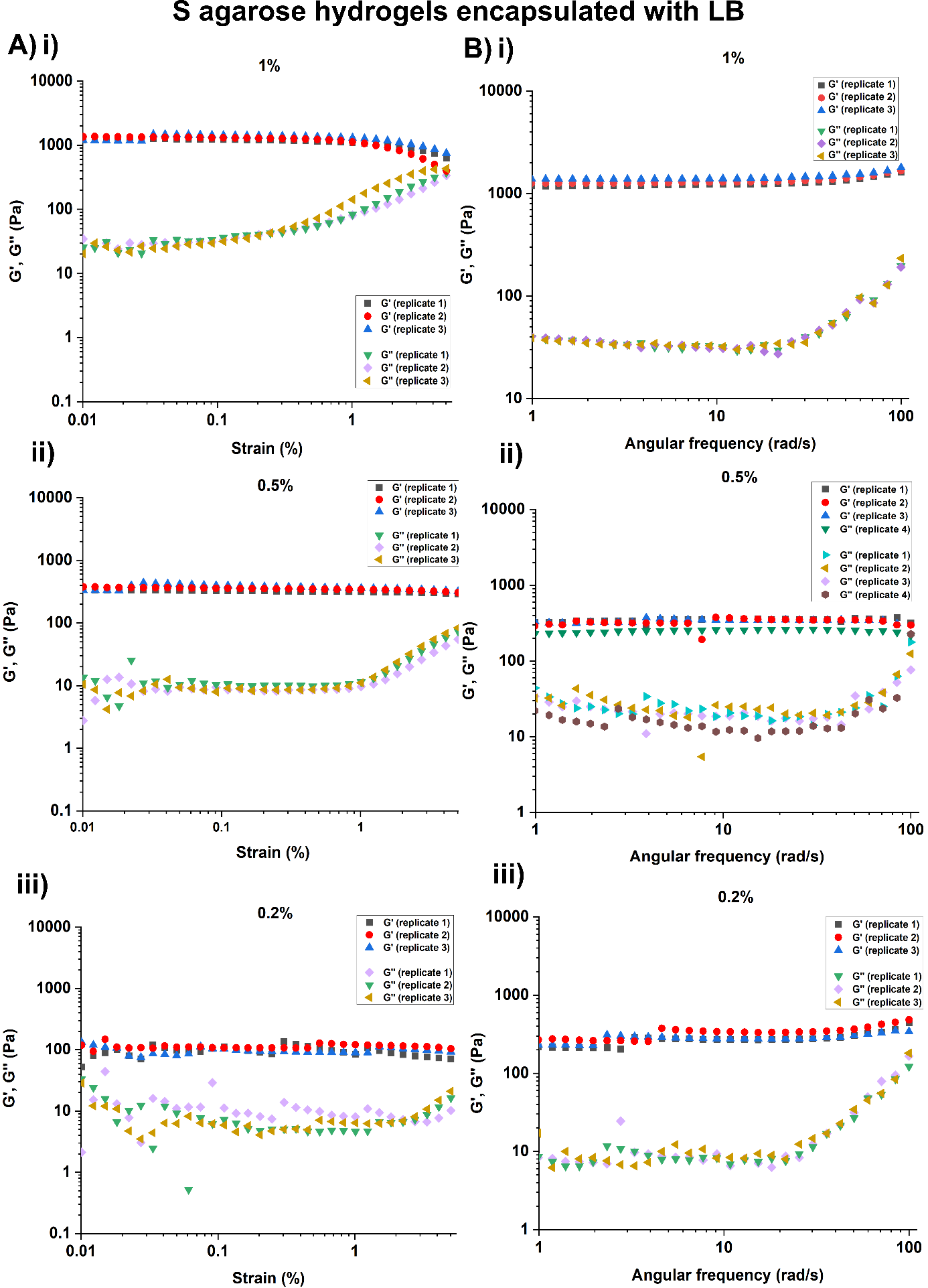

**Figure S17.** Rheology analysis. (A) Amplitude and (B) frequency sweeps of substituted agarose hydrogels encapsulated with LB. All experiments were performed in three replicates – (i), (ii), (iii).

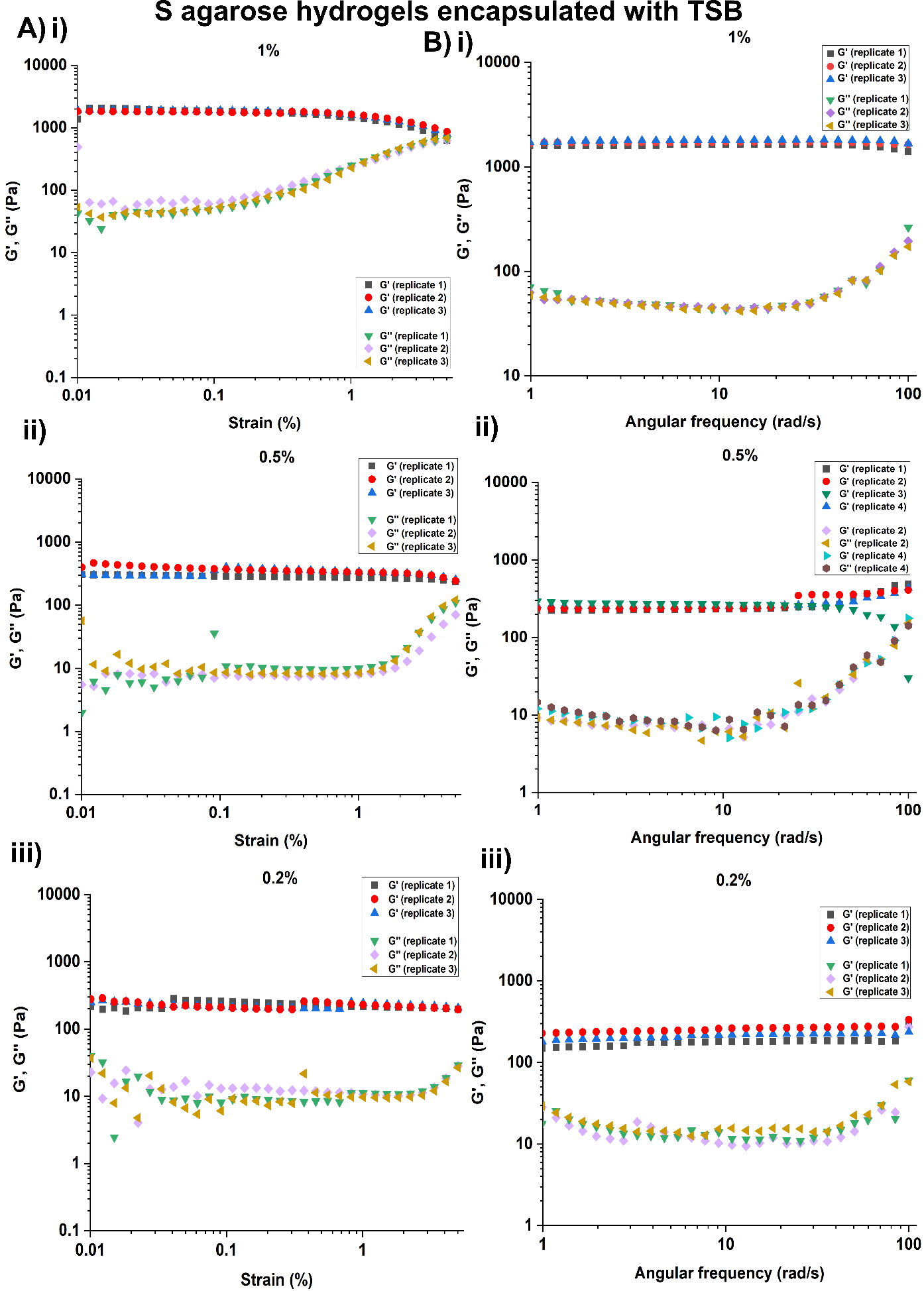

**Figure S18.** Rheology analysis. (A) Amplitude and (B) frequency sweeps of substituted agarose hydrogels encapsulated with TSB. All experiments were performed in three replicates – (i), (ii), (iii).

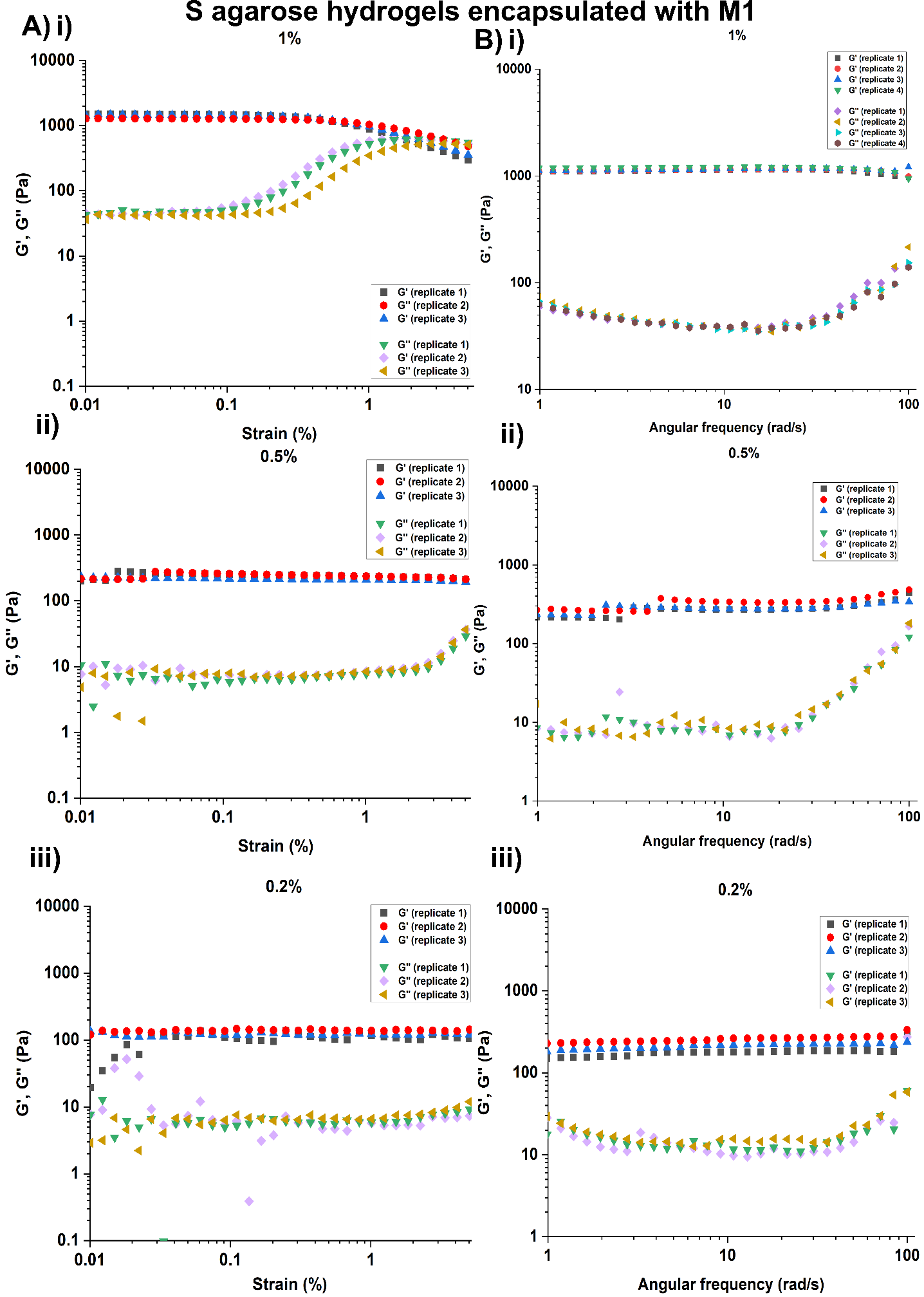

**Figure S19.** Rheology analysis. (A) Amplitude and (B) frequency sweeps of substituted agarose hydrogels encapsulated with M1. All experiments were performed in three replicates – (i), (ii), (iii).

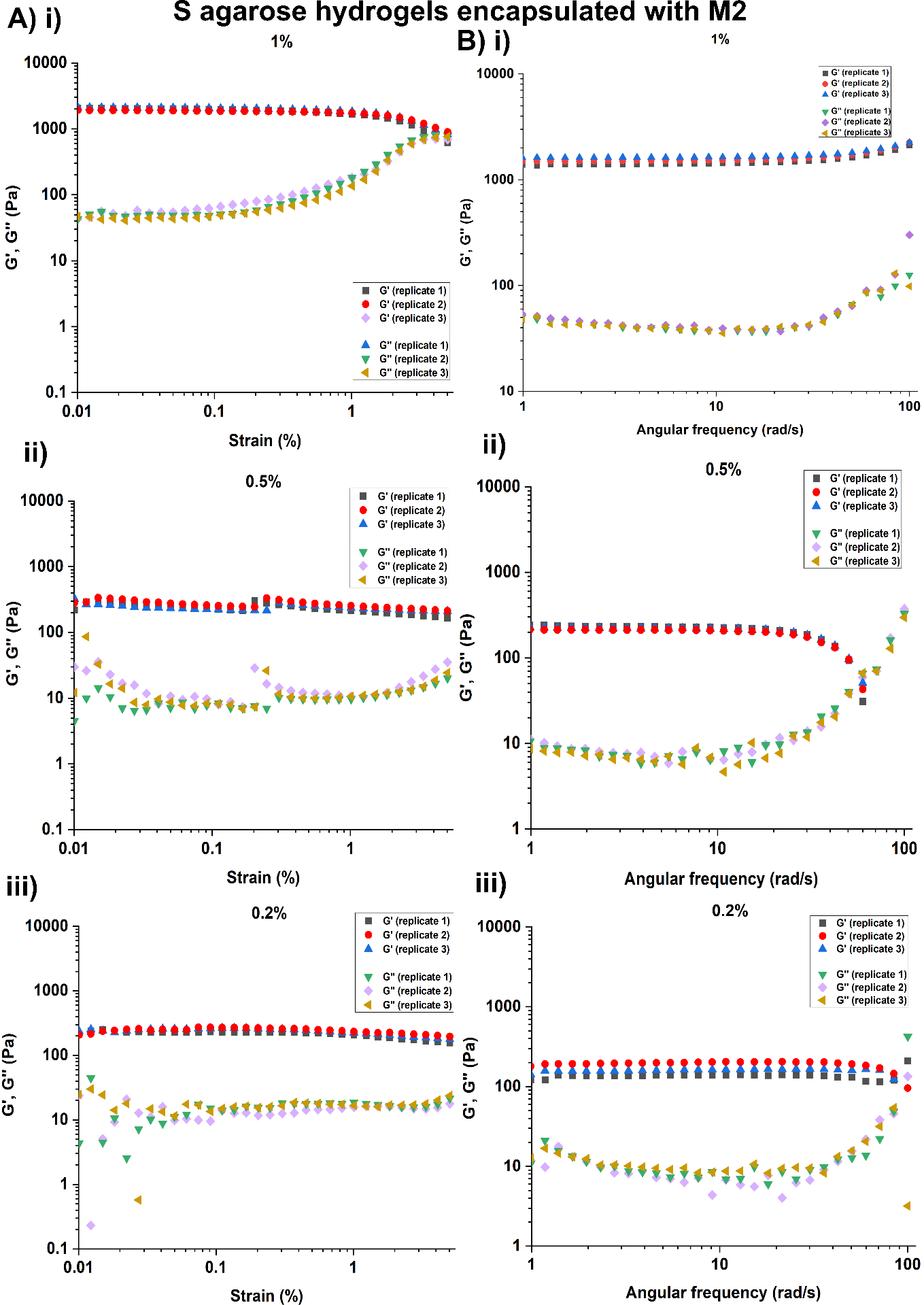

**Figure S20.** Rheology analysis. (A) Amplitude and (B) frequency sweeps of substituted agarose hydrogels encapsulated with M2. All experiments were performed in three replicates – (i), (ii), (iii).

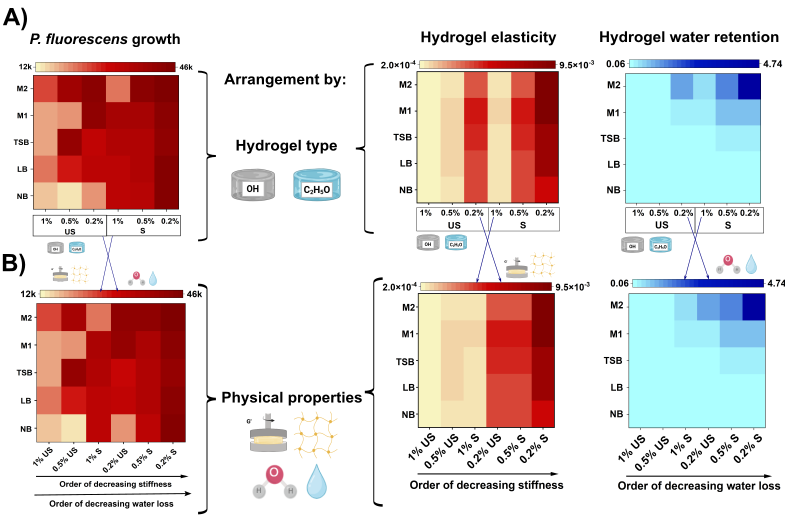

**Figure S21.** Heatmaps indicating the growth of *P. fluorescens* according to **(A)** hydrogel type, hydrogel elasticity and water retention and **(B)** in order of decreasing hydrogel stiffness and water loss.

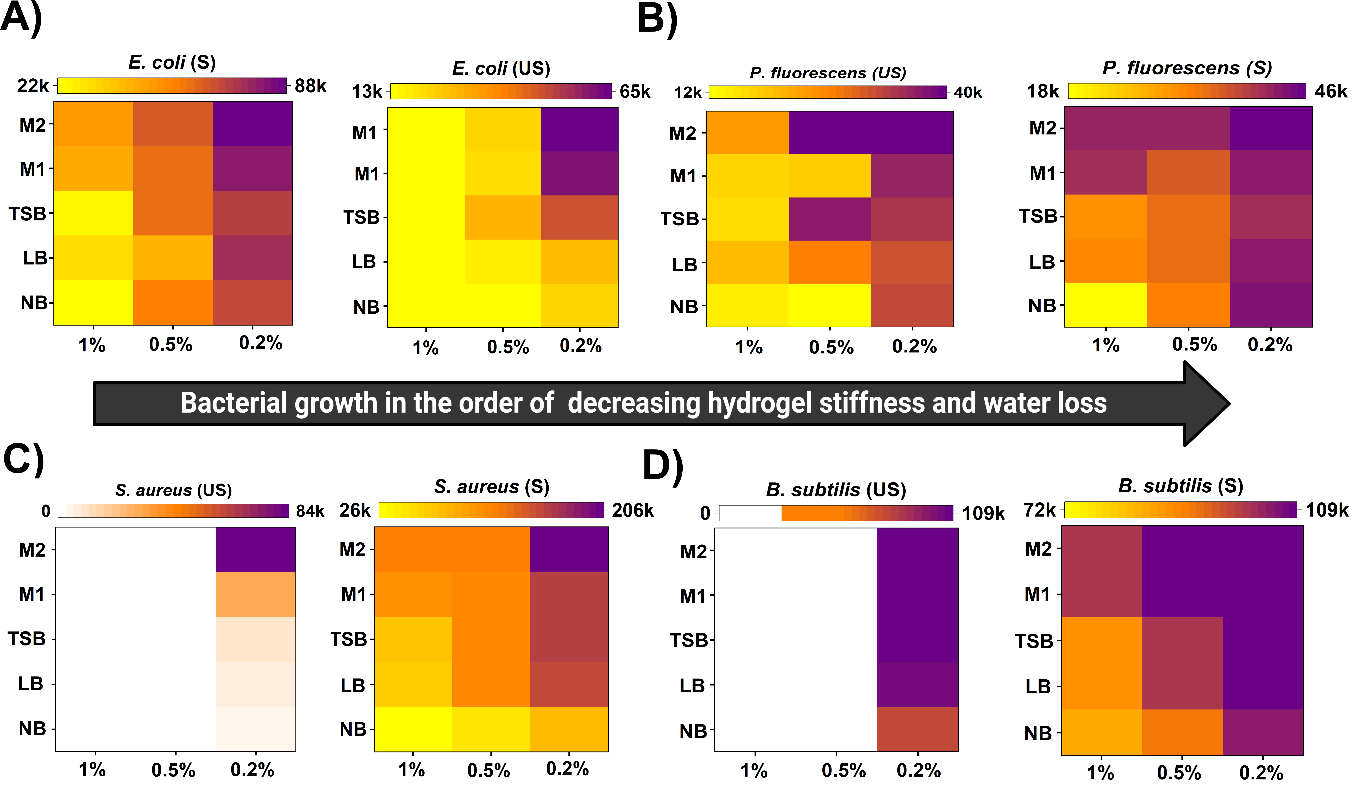

**Figure S22.** Heat maps of **(A)** *E. coli*, **(B)** *P. fluorescens*, **(C)** *S. aureus*, and **(D)** *B. subtilis* growth across hydrogel types and concentrations, aligned with decreasing stiffness and water loss.

|  | Correlation analysis | |
| --- | --- | --- |
| Bacterial species | Bacterial growth and  hydrogel stiffness | Bacterial growth and hydrogel water loss |
| *E. coli* | -0.804 (***) | -0.648 (***) |
| *P. fluorescens* | -0.664 (***) | -0.564 (***) |
| *S. aureus* | -0.663 (***) | -0.615 (***) |
| *B. subtilis* | -0.751 (***) | -0.632 (***) |

**Table S23.** Spearman’s correlation co-efficient indicating negative correlation between bacterial growth and hydrogel stiffness, and hydrogel water loss.

| Bacterial species | Agarose hydrogel type | Correlation co-efficient  (bacterial growth and hydrogel stiffness) | Correlation co-efficient (bacterial growth and hydrogel water loss) |
| --- | --- | --- | --- |
| *E. coli* | US | -0.851 (***) | -0.758 (***) |
|  | S | -0.725 (***) | -0.502 (***) |
| *P. fluorescens* | US | -0.537 (***) | -0.78 (***) |
|  | S | -0.562 (***) | -0.302 (***) |
| *S. aureus* | US | -0.817 (***) | -0.601 (***) |
|  | S | -0.635 (***) | -0.857 (***) |
| *B. subtilis* | US | -0.823 (***) | -0.592 (***) |
|  | S | -0.545 (***) | -0.676 (***) |

**Table S24**. Spearman’s correlation co-efficient indicating negative correlation between bacterial growth and hydrogel stiffness, and hydrogel water loss for US and S agarose hydrogels.

|  | 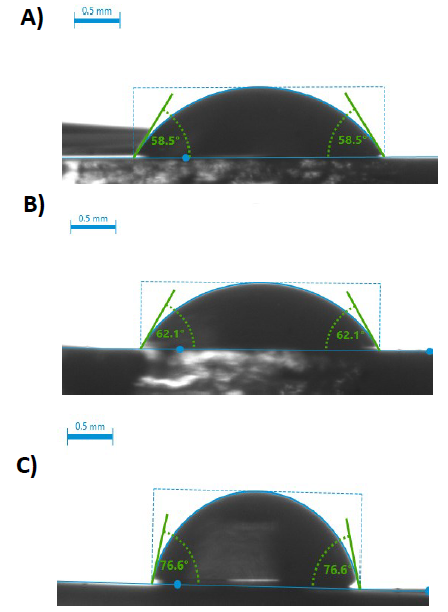 | | | |
| --- | --- | --- | --- | --- |
|  | **Figure S25.** Contact angle analysis of substituted hydrogels indicating hydrophobicity of hydrogels. Contact angle of (A) 1% (mean value = 58.5°), (B) 0.5% (mean value = 62.1°), and 0.2% (mean value = 76.6°) unsubstituted hydrogels with water drop (volume = 5 μL). | | | |
| Effect | | Mean Square | F-statistic | p-value  (Pr (>F)) |
| Main Effects | |  |  |  |
| Hydrogel concentration | | 5.024e+10 | 814.091 | < 2e-16 *** |
| Bacterial species | | 2.913e+10 | 471.975 | < 2e-16 *** |
| Hydrogel type | | 2.354e+11 | 3814.212 | < 2e-16 *** |
| Nutrient Media | | 9.363e+09 | 151.724 | < 2e-16 *** |
| Two-way Interactions  (Factor variable 1: Factor variable 2) | |  |  |  |
| Hydrogel concentration: Bacterial species | | 5.887e+09 | 95.397 | < 2e-16 *** |
| Hydrogel concentration: Hydrogel type | | 6.641e+09 | 107.619 | < 2e-16 *** |
| Hydrogel concentration: Nutrient media | | 8.401e+08 | 13.614 | 3.07e-16 *** |
| Bacterial species: Hydrogel type | | 3.811e+10 | 617.538 | < 2e-16 *** |
| Bacterial species: Nutrient media | | 1.841e+09 | 29.825 | < 2e-16 *** |
| Hydrogel type: Nutrient media | | 6.800e+08 | 11.019 | 3.20e-08 *** |
| Three-way Interactions  (Factor variable 1: Factor variable 2: Factor variable 3) | |  |  |  |
| Hydrogel concentration: Bacterial species: Hydrogel type | | 7.610e+09 | 123.312 | < 2e-16 *** |
| Hydrogel concentration: Bacterial species: Nutrient media | | 1.567e+08 | 2.540 | 0.000182 *** |
| Hydrogel concentration: Hydrogel type: Nutrient media | | 1.186e+09 | 19.220 | < 2e-16 *** |
| Bacterial species: Hydrogel type: Nutrient media | | 1.738e+09 | 28.167 | < 2e-16 *** |
| Four-way Interaction  (Factor variable 1: Factor variable 2: Factor variable 3: Factor variable 4) | |  |  |  |
| Hydrogel concentration: Bacterial species: Hydrogel type: Nutrient media | | 3.398e+08 | 5.507 | 5.88e-13 *** |

**Table S26.** Four-way ANOVA analysis for determining the impact of hydrogel properties on bacterial growth. p-values: * = 0.05, ** = 0.01, *** = 0.001.

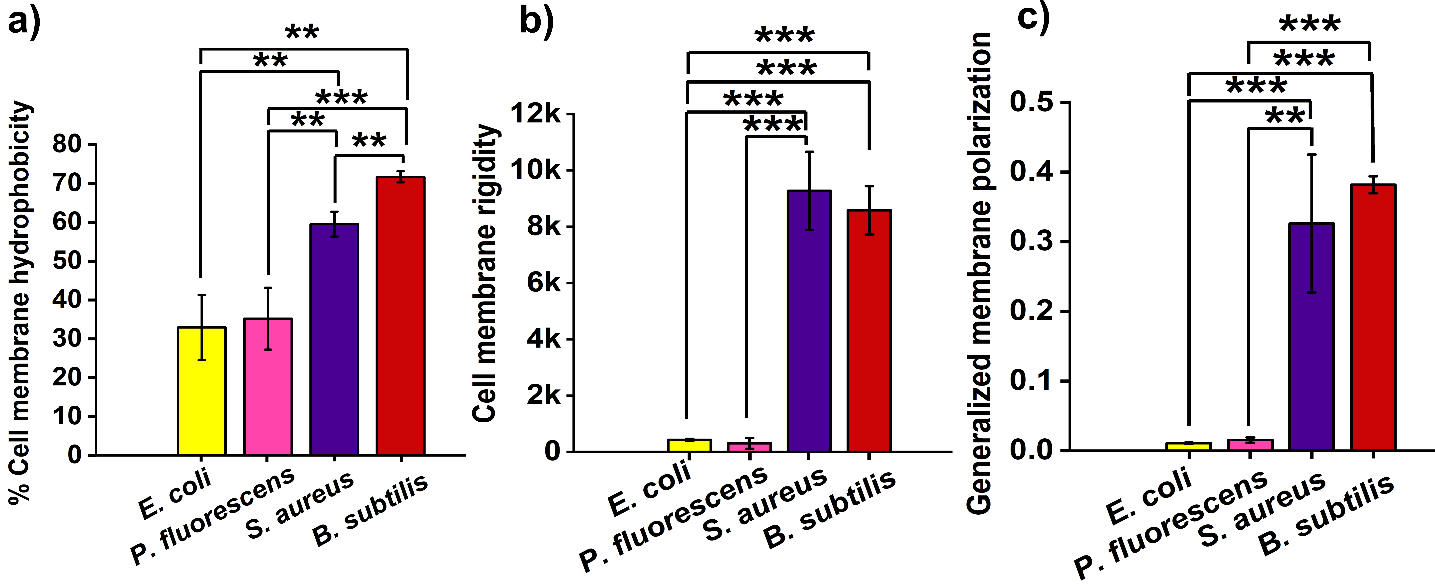

**Figure S27**. Bacterial cell membrane characterisitics. a) cell membrane hydrophobicity, b) cell membrane rigidity, and c) membrane polarization. p-values: * = 0.05, ** = 0.01, *** = 0.001. Across all the tested cell membrane properties, *S. aureus* and *B. subtilis* demonstrated increased hydrophobicity, rigidity and polarization compared to *E. coli* and *P. fluorescens*.

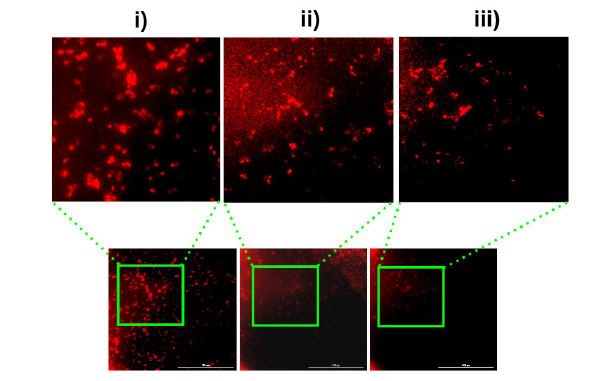

**Figure S28.** Propidium iodide staining of *S. aureus* cells on 1% US agarose hydrogels across 3 replicates (i), (ii), and (iii) indicating compromise or damage of bacterial cell membranes. To confirm if bacteria are dead, the hydrogels were cut and transferred into resazurin solution to check for *S. aureus* viability.

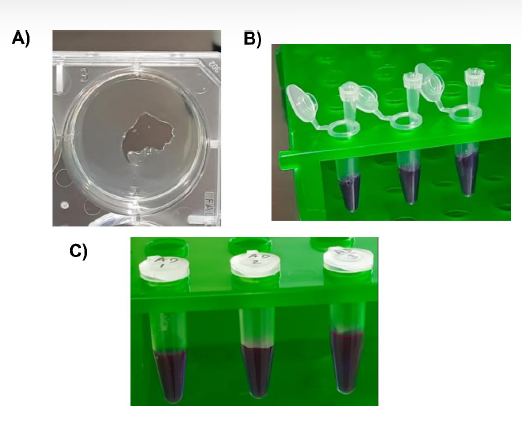

**Figure S29.** *S. aureus* viability confirmation by end point resazurin assay. A) A portion of agarose hydrogel where *S. aureus* was introduced was excised and transferred into B) resazurin solution. C) End point viability result after 18 hours incubation at 37°C indicating non-viability of cells as seen by the lack of colour change to pink.

### **S30. Cell membrane hydrophobicity - Microbial adhesion to hydrocarbons (MATH) test**

n-hexane hydrocarbon was used for determining the cell membrane hydrophobicity. 4 mL of bacterial culture OD_600nm_ ~ 0.5 – 0.7 was mixed with 1 mL of n-hexane solution. The mixture was thoroughly vortexed for 2 minutes and allowed to phase separate at room temperature for 15 minutes. Hydrophobic bacteria will be bound to n-hexane hydrocarbon molecules and travel to the top layer of the tube. This solution appears yellow brown in colour. In contrast, hydrophilic bacteria will be present in the clear and transparent bottom layer of the tube. 750 µL of the solution from bottom layer (test solution) is used for measurement at 400 nm – 600 nm. Bacterial culture media ((LB for *E. coli*, *P. fluorescens*, and *B. subtilis*) and (TSB for *S. aureus*)) without any bacteria was used as blank for the respective bacterial species. % Cell surface hydrophobicity was calculated by the following formula:

$$\% Cell surface hydrophobicity=$$

$100 x \frac{\left( Absorbance of control-Absorbance of test solution \right)}{Absorbance of control}$

### **S31. Cell membrane rigidity – Nile red assay**

The rigidity of bacterial cell membranes was evaluated using the Nile red, a fluorescent (excitation wavelength: ~ 530-550 nm and emission wavelength: ~ 590-700nm, depending on the solvent) dye which binds to the lipid groups present in the membranes. The dye was obtained from the School of Life Sciences, University of Warwick. Bacteria from primary culture (section 3.3.1) were sub-cultured and diluted to OD_600nm_ ~ 0.5 – 0.7. The cultures were washed and suspended in 1XPBS. The cultures were then introduced in three replicates in 96-well black polystyrene plate and incubated with 2µg/mL Nile red dye for 10 minutes at room temperature under dark conditions. Post-incubation, kinetic fluorescence measurements were recorded from 400 nm to 700 nm to determine the peak Nile red intensities across bacterial species.

### **S32. Cell membrane polarization - Laurdan assay**

The polarity of bacterial cell membranes was determined using Laurdan, a fluorescent dye (excitation wavelength: 350 nm and emission wavelength: 460 nm and 500 nm) purchased from Sigma Aldrich, Merck. The dye intercalates into bacterial cell membranes and produces a shift in its wavelengths depending on the number of water molecules present in the membrane. Bacteria from primary culture (section 3.3.1) were sub-cultured and diluted to OD_600nm_ ~ 0.5 – 0.7. These bacterial cultures were then incubated with 100 µM Laurdan for 5 minutes at room temperature under dark conditions. Post-incubation, the samples were washed four time with PBS to remove membrane un-interacted Laurdan dye molecules. Post-washing, 200 µL of the sample was transferred into sterile 96-well black polystyrene plate. The experiments were performed in three replicates for each bacterial species. Membrane polarization was determined by measuring the fluorescence at 460 nm and 500 nm using BioTek Cytation 5 Cell Imaging Multimode Reader (Agilent).

Generalized membrane polarization was calculated by:

$$emission intensity at 460 nm-emission intensity at 500 nm$$

$emission intensity at 460 nm+emission intensity at 500 nm$
